# Self-organized and self-sustained ensemble activity patterns in simulation of mouse primary motor cortex

**DOI:** 10.1101/2025.01.13.632866

**Authors:** D.W. Doherty, J. Jung, Dura-Bernal, W.W. Lytton

## Abstract

The idea of self-organized signal processing in the cerebral cortex has become a focus of research since Beggs and Plentz ^1^ reported avalanches in local field potential recordings from organotypic cultures and acute slices of rat somatosensory cortex. How the cortex intrinsically organizes signals remains unknown. A current hypothesis was proposed by the condensed matter physicists Bak, Tang, and Wiesenfeld ^2^ when they conjectured that if neuronal avalanche activity followed inverse power law distributions, then brain activity may be set around phase transitions within self-organized signals. We asked if we would observe self-organized signals in an isolated slice of our data driven detailed simulation of the mouse primary motor cortex? If we did, would we observe avalanches with power-law distributions in size and duration and what would they look like? Our results demonstrate that a brief unstructured stimulus (100ms, 57*μ*A current) to a small subset of neurons (about 181 of more than 10,000) in a simulated mouse primary motor cortex slice results in self-organized and self-sustained avalanches with power-law size and duration distributions and values similar to those reported from in vivo and in vitro experiments. We observed 4 cross-layer and cross-neuron population patterns, 3 of which displayed a dominant rhythmic component. Avalanches were each composed of one or more of the 4 population patterns.

## Introduction

A promising approach to understanding signal processing among neurons in the cerebral cortex has been inspired by condensed matter physics where, starting from well characterized individual components, a rigorous framework has been constructed to explain and predict properties that appear due to component interactions ^2,3^. Beggs and Plentz ^1^ pioneered investigating neuronal avalanche activity postulated by Bak, Tang, and Wiesenfeld ^2^ using local field potential recordings from organotypic cultures and acute slices of rat somatosensory cortex. Since their original report, Beggs, Plenz, and many others have investigated neuronal avalanche activity in vitro ^4–7^ and in vivo ^8–18^. The observation of avalanches with long-tailed size and duration distributions is consistent with the idea that cortical activity may be set around phase transitions within self-organized signals known as a critical cortical state or criticality ^2^.

Accumulating experimental evidence is consistent with the hypothesis that signal processing in the cerebral cortex is maintained near criticality ^12,13,19–23^. Synchronized activity among cortical neurons spans a broad range of population sizes and durations at criticality. In contrast, activity with little to no synchronized responses is known as subcritical activity and a substantial population of neurons with synchronized activity is supercritical activity ^24,25^. Experimental evidence shows that cortical activity consistent with criticality may be pushed into highly active synchronous spiking by suppressing inhibition using GABA antagonists. In contrast, these same neurons may be pushed into low activity asynchronous spiking by increasing inhibition with GABA agonists or decreasing excitation with AMPA and NMDA antagonists ^26–30^. These data suggest that a proper balance must be maintained between excitation and inhibition for the cortex to carry out signal processing near criticality ^31–34^. Recent evidence shows that, in addition to the balance between inhibition and excitation, synaptic strength may play an important part in criticality ^35^. Cortical activity that communicates through weak synapses may operate at criticality without any inhibition. With weak synapses criticality is strong and is associated with large fluctuations in activity. But it is fragile. Adding just a small amount of inhibition moves the population away from criticality. A population of cortical neurons with strong synapses results in weaker but broad robust criticality with smaller fluctuations in activity.

A major challenge in determining if brain structures generate self-organized signals and what those signals look like is the inability to record from every neuron in a structure. In mammalian brains the number of cells recorded from has been far fewer than the number comprising the structures of interest. This is known as the subsampling problem ^10,36^. Two-photon calcium imaging is currently helping to close the gap on numbers of neurons concurrently recorded from a brain structure and the structure’s neuronal population ^37–40^. Indeed, some of these studies have described self-organized emergent signals through avalanche activity and other measures ^38,41^. Two-photon calcium imaging has even been used to simultaneously record from every neuron in the zebrafish larva brain ^42^. The research team concluded that zebrafish larva brain activity is organized into scale-invariant neuronal avalanches.

However, two-photon calcium imaging remains unable to record from every neuron, or even a majority of neurons in mammalian brain structures. Perhaps even more importantly, one is unable to assign specific morphology and connectivity to each recorded neuron. Using a detailed simulation of the mouse primary motor cortex (M1) we set out to understand the relationship between power-law activity patterns, their values, and the neural responses observed from every neuron across different layers, cell populations, and the entire cortical column.

## Results

For this study, 490 simulations of durations of 1-10 min simulation time were run over >1M core hours of CPU time requiring over >1M core hours of CPU time. A typical 10 min simulation took ∼100 hours to complete across 192 cores of high performance computer (HPC) running Intel Skylake CPUs using a fixed timestep of 0.05 ms under NetPyNE/NEURON.

### Sustained oscillations with no ongoing input

A brief, focal stimulation (100 ms; 0.57 nA/cell, 1.5%) produced continuing population bursting activity for as long as the recording continued (10 min) in 7 networks with different random wiring. A typical data set (Fig 1) began with sparse stimulation activation (Fig 1B left) followed by a ∼500 ms narrow band of sustained deep layer activity (up to first asterisk in Fig 1A). This activity then increased dramatically in the deep layers and spread to all layers, producing a characteristic recurring pattern occurring at about 1 Hz. Each instance of this *delta pattern* followed this motif of sparse activity in deep layers followed by increasingly intense activation with spread (Fig 1A between row 1 asterisks; Fig 1B).

**Figure 1.**
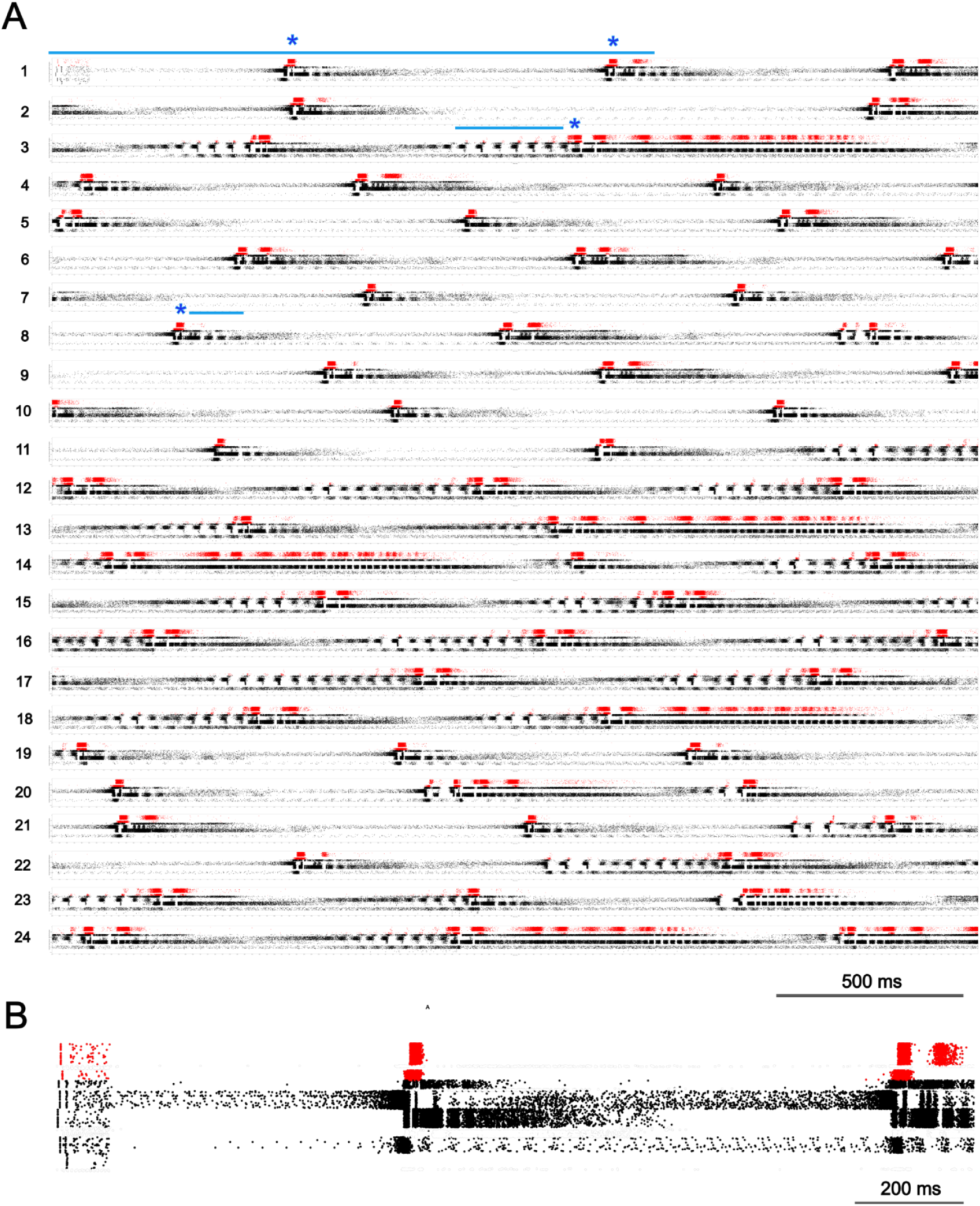
Raster plot of excitatory neuron firing in the first 1 minute of a 10 min M1 simulation. (A) Continuous (wrapped) 60 s raster plot (24 rows of 2.5 s each; red: L2-4; black: L5-6) Sustained activity was initiated by a brief stimulus (100 ms; 0.57 nA/cell, 1.8% of cells; top left). Asterisks denote the start of selected delta events. Row 1 line is expanded in B; row 3 line above beta event; row 8 line above gamma event. (B) Detail of first 1.64s (row 1 blue line in A).

The entire 10 min simulation was characterized by the repeated cross-layer bursts shown in Fig 1A. The prolonged delta pattern bursts produced activity of varying duration and some displayed multiple bursts of superficial layer activity -- note the repeat of red in some of the events (Fig 1A). Compare the first delta event in Fig 1A row 1, which was about 400 ms in duration (left, blue asterisk) and displayed a single burst of activity in superficial layers (red rasters), and the delta event in the right half of row 3 (Fig 1A blue asterisk row 3), which was about 1 s in duration and had at least 10 bursts within superficial layers. Delta bursts were similar in activity pattern across time in individual simulations and across different simulations (6 networks). Delta patterns began with intense activity in 2 narrow bands within deep layers (Fig 1B), followed by intense activity between the 2 bands and above the most superficial band. Intense activity then appeared in superficial layers (Fig 1B, red).

The delta pattern was frequently preceded by multiple brief bursts of cross-layer activity (Fig 1; blue line left of blue asterisk row 3), at ∼22 Hz, beta frequency. This beta pattern began with rapid increase in a narrow band of deep layer activity but, in contrast with the onset of the delta pattern, activity that started just superficial to the narrow band, increased at a slow rate and peaked towards the end of a single period. Another activity pattern was in the gamma range -- ∼54 Hz. This gamma pattern displayed ∼18.5 ms recurrent bursts of strong activity in a narrow band within deep layers, nested in qualitatively identified delta events (Fig 1A, row 8 blue line following a blue asterisk). Rasters were rarely seen in a band above the periodically active band during gamma.

We set out to quantitatively assess the consistency of the 3 frequency dominated delta, beta, and gamma patterns across time and datasets (Fig 2). To further investigate each of the recurrent patterns associated with dominant frequencies, we used a time-domain features algorithm known as SPUD ^43^. Using the SPUD software, we quantitatively assessed the consistency of the delta pattern (Fig 2A) across time and datasets. Delta patterns were selected from large IT2/3 activity peaks (>= 250 spikes) from 1 ms binned histograms of cortical column activity. IT2/3 peaks less than or equal to 50 ms apart were filtered out. Activity that expanded beyond layers 5B and 6 (IT5B/IT6) displayed approximately one delta pattern of activity every second.

**Fig 2.**
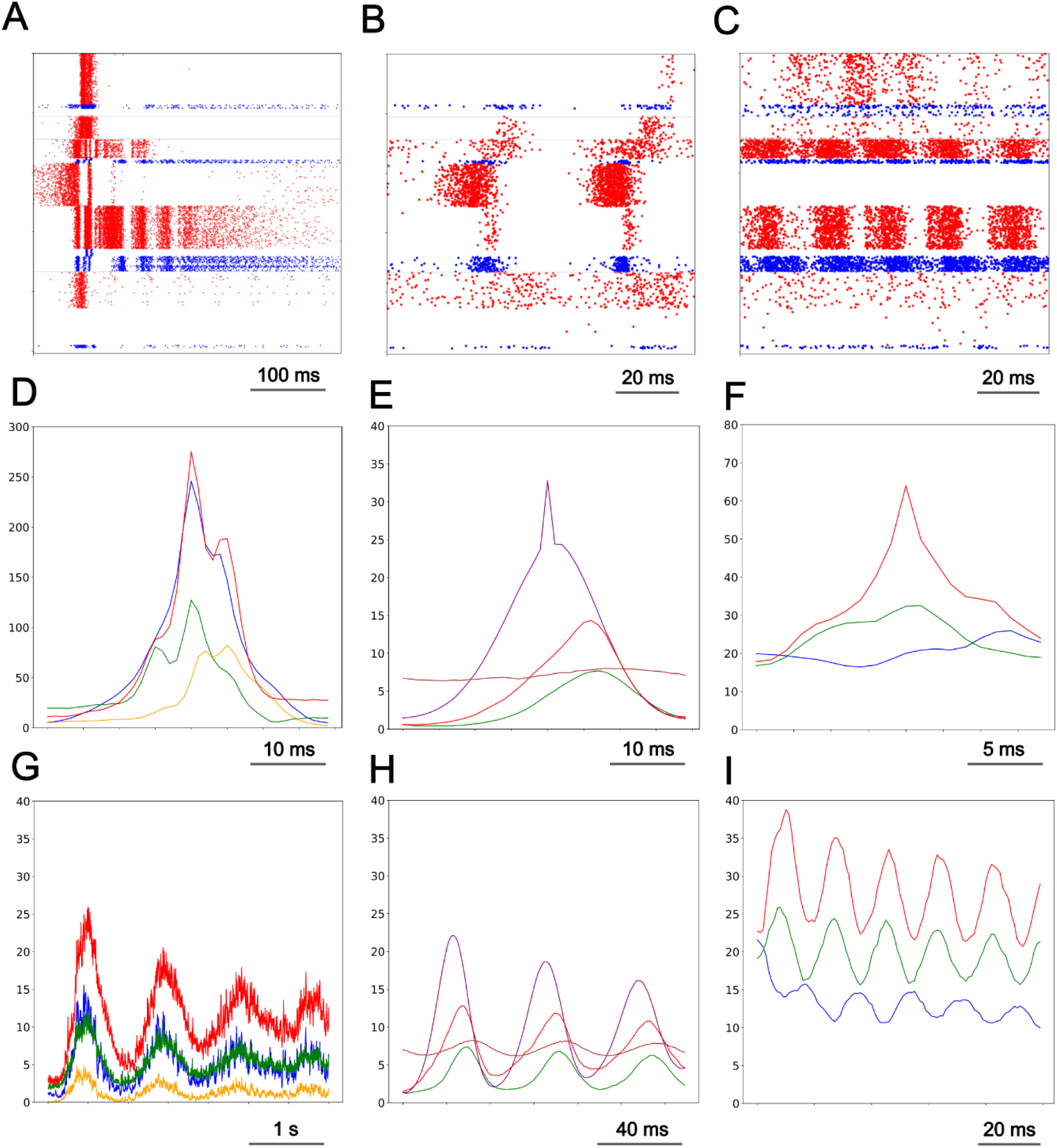
Recurring population events. A-C. Raster plots for delta, beta, gamma patterns respectively; red: excitatory; blue: inhibitory cells). D-F. Average spike histograms showing 4 periods of delta, 3 of beta, 3 of gamma (note different time scales) G-I. Characteristic initiation for each pattern. (in D-I -- IT2/3:blue; IT4:yellow; IT5A:green; IT5B:purple; PT5B:red; IT6:brown)

Automated pattern selection identified 1,878 individual delta patterns measured across 6 datasets (Fig 2D). The delta pattern activity peaks were characterized as concurrent IT2/3 and PT5B activity peaks (Fig 2D blue and red respectively) with a rise in IT5A activity during onset (Fig. 2D green) followed by a rise in IT4 activity during the decrease in IT2/3 and PT5B activity from their initial activity peaks (Fig. 2D yellow). The end of each delta pattern envelope was marked by ongoing PT5B neuron activity (Fig 2ADG, red), notable because PT5B is the primary extra-telencephalic outflow pathway.

We used SPUD ^43^ analysis to quantitatively assess the consistency of the beta pattern (Fig 2B) across time and datasets. Unique to the beta pattern was that spiking activity was visible in layer 2/3 inhibitory but not excitatory neurons (Fig 2B; superficial blue rasters). First SPUD software was used to pick IT5B peak activity (>= 15 spikes) from cortical activity, which found all delta and beta pattern activity. Next, IT5B activity that occurred during robust IT2/3 activity (>= 250 spikes), activity unique to the delta pattern, was filtered out. Automated pattern selection identified 11,163 periods of beta pattern activity across 6 statistically unique data sets.

Beta wave phases did not match up when averaged across dataset histograms. Therefore, we selected the single dataset with the largest number of beta pattern periods, 6,909, and provided the average histograms for participating neuron populations (Fig. 2EH). Beta pattern IT5B activity was most prominent (Fig. 2EH purple) followed by PT5B (red) and IT5A activity (green). Ongoing IT6 neuron activity (brown) increased and decreased with IT5A activity. Beta patterns, when present, were seen preceding the onset of delta patterns (Fig. 1A, 4A).

Quantitative assessment of the gamma pattern (Fig 2C) consistency was carried out using the SPUD software package. Gamma pattern excitatory neurons IT2/3, IT5A, and PT5B displayed clearly periodic patterns as did the inhibitory PV5B neurons (Fig 2C). The software picked PT5B peak activity (>= 20 spikes), which is prominent in the gamma pattern but also in the delta pattern. Next, IT2/3 activity that occurred during robust PT5B activity (>= 250 spikes) was filtered out. This last step removed delta pattern activity. Automated pattern selection found 2,945 periods of gamma pattern across 6 statistically unique data sets. Gamma wave phases did not match up when we averaged across dataset histograms. Therefore, we selected the single dataset with 503 gamma waves and provided the average histograms for participating neuron populations (Fig 2FI). The gamma pattern displayed ∼54 Hz oscillation most strikingly in PT5B and IT5A (Fig 2FI, red and green). Gamma oscillations also appeared in IT2/3 (Fig 2F, blue), which were anticorrelated with IT5A and PT5B activity (Fig 2I). Prominent IT5B activity was mostly absent during gamma pattern activity, in contrast with during delta or beta pattern activity.

We qualitatively and quantitatively identified delta, beta, and gamma rhythm dominated patterns in 6 of 7 simulation configurations, all with identical wiring densities between neuronal populations but unique connectivity defined using random seeds. Spectrograms demonstrated power in the delta range and across-frequency power (Fig. 3C; top and middle) aligned with each 1 Hz burst of spiking activity in the raster plot below (Fig. 3C; bottom), confirming the delta rhythm is associated with the delta pattern. Likewise, spectrograms demonstrated power in the beta range associated with beta pattern activity (Fig 4C) and power in the gamma range nested within the delta pattern (Fig 4B). A fourth pattern was identified, the irregular pattern, that was composed exclusively of IT5B and / or IT6 activity and was not rhythm dominated. One of the 7 simulation configurations displayed sustained irregular pattern activity for the duration of one 10 minute simulation.

**Fig 3.**
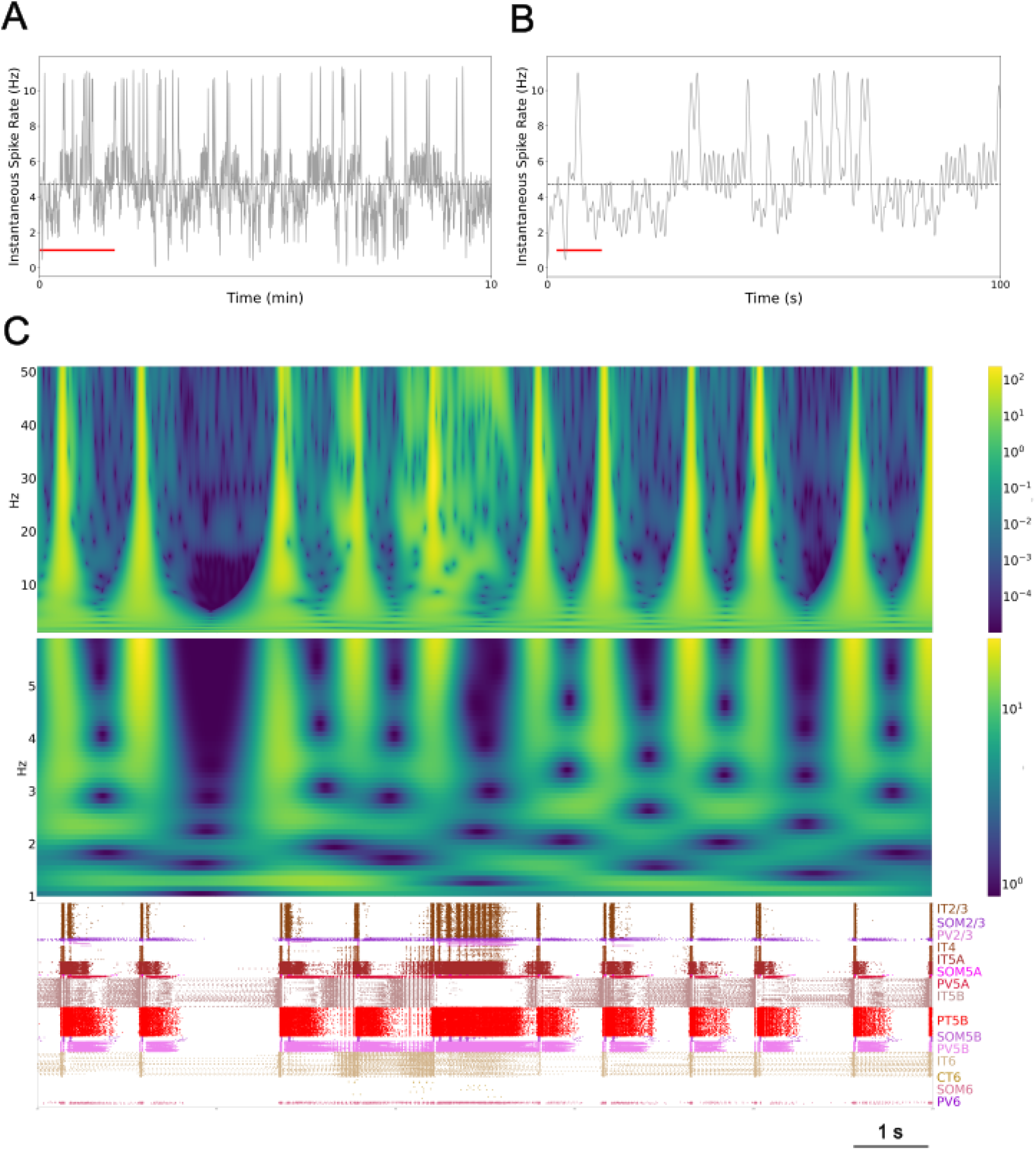
Self-sustained activity showed oscillatory ∼1 Hz activity. (A) Instantaneous spike rates over 10 min, all neurons; mean firing rate is 4.7 Hz (horizontal line); CV 1.53. (B) Detail of first 100 s above red line in A. (C) Spectrograms (top 0-50 Hz; middle 0-5 Hz) and raster (10 s, red line in B; population labels at right).

**Fig 4.**
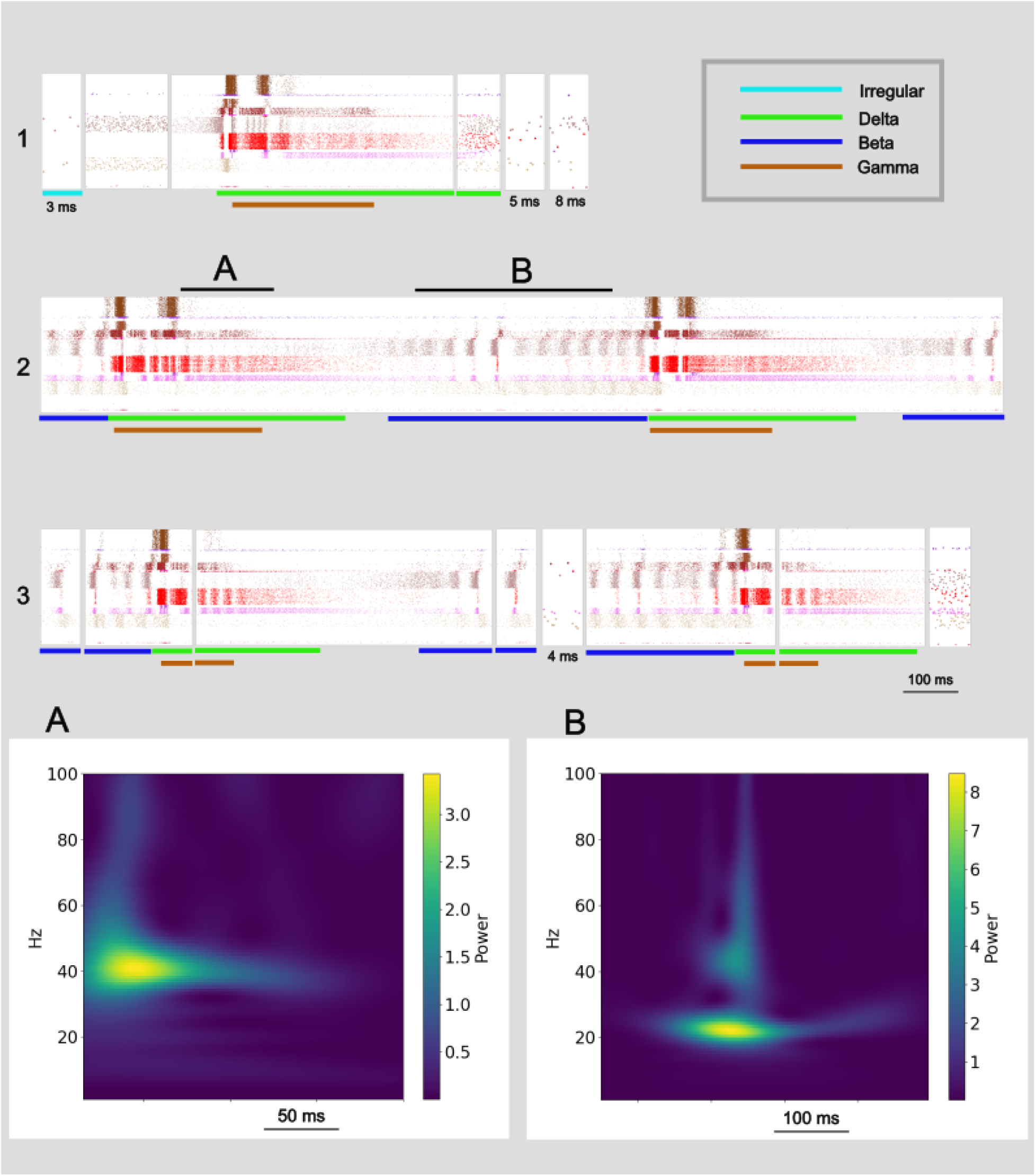
Oscillatory activity patterns make up individual avalanches. Continuous sequences of raster plots divided into individual avalanches. Top row is one continuous sequence. Bottom two rows compose another continuous sequence. **A, B.** Spike rate spectrograms of segments marked above show strong gamma activity, beta activity, respectively.

We have demonstrated that the self-organized and self-sustained pattern of activity in data-driven simulated cortical slices are composed of 3 rhythm dominant patterns and 1 irregular pattern. Cortical activity patterns alternate between delta patterns preceded by few to no visible beta events as seen in Fig 1 rows 4-10 (beta sparse) and delta events preceded by many beta events (Fig 1 rows 12-18; beta rich). A spike rate histogram of the entire 10 minute simulation (Fig. 3A) showed periods of low activity (∼3.2 Hz) interspersed with periods of high activity (∼6.0 Hz; overall 4.7 Hz average shown as dashed line) and an infrequent activity level that peaked around 11 Hz. Low ∼3.2 Hz instantaneous spike rates are aligned with beta sparse delta activity displayed at left and right in the Fig 3C raster plot (bottom; Fig 3B red line). In contrast, high ∼6.0 Hz spike rates are aligned with beta rich activity (Fig 3C, center left). The 11 Hz peak in instantaneous spike rate (Fig. 3B; middle above red line) is associated with higher power in the gamma range (Fig. 3C; top) and is aligned with extended nested gamma pattern activity displayed in the raster plot (Fig. 3C; bottom).

### Avalanche power-law distributions

Next we assessed the self-organized and self-sustained activity driven by a brief stimulus to our simulated M1 cortical slice for clusters of spiking activity with distributions typical of avalanche activity. We placed all simulated spiking activity into 1 millisecond bins and defined an avalanche as composed of adjacent bins filled with one or more action potentials, preceded and followed by at least one empty bin. The total number of action potentials in the filled bins equaled the avalanche size and the product of the total number of contiguously filled bins and bin duration (1 ms) equaled the avalanche duration. We observed 15,578 avalanches in our representative 10 minute simulation (Fig. 1).

Self-sustained responses with consistent expressions of bursts did indeed fit power-law size distributions in the range of −1.5 but fit duration distributions around −1.6 rather than −2.0. The pair of log-log graphs in Fig 5AB display the distribution of avalanches from our representative 10 minute simulation of M1 activity (Fig. 1). Self-sustained responses without consistent expressions of bursts of activity across layers and neuron cell types did not fit power-law distributions. Those active in only IT5B/IT6 (irregular pattern) were composed of small and short duration avalanches and, therefore, lacked long tailed avalanche distributions. Sustained activity with some bursts of activity across layers resulted in increases in the number of midsize and large avalanches in proportion to the number of bursts, but not enough for robust fits to power-law distributions.

**Figure 5.**
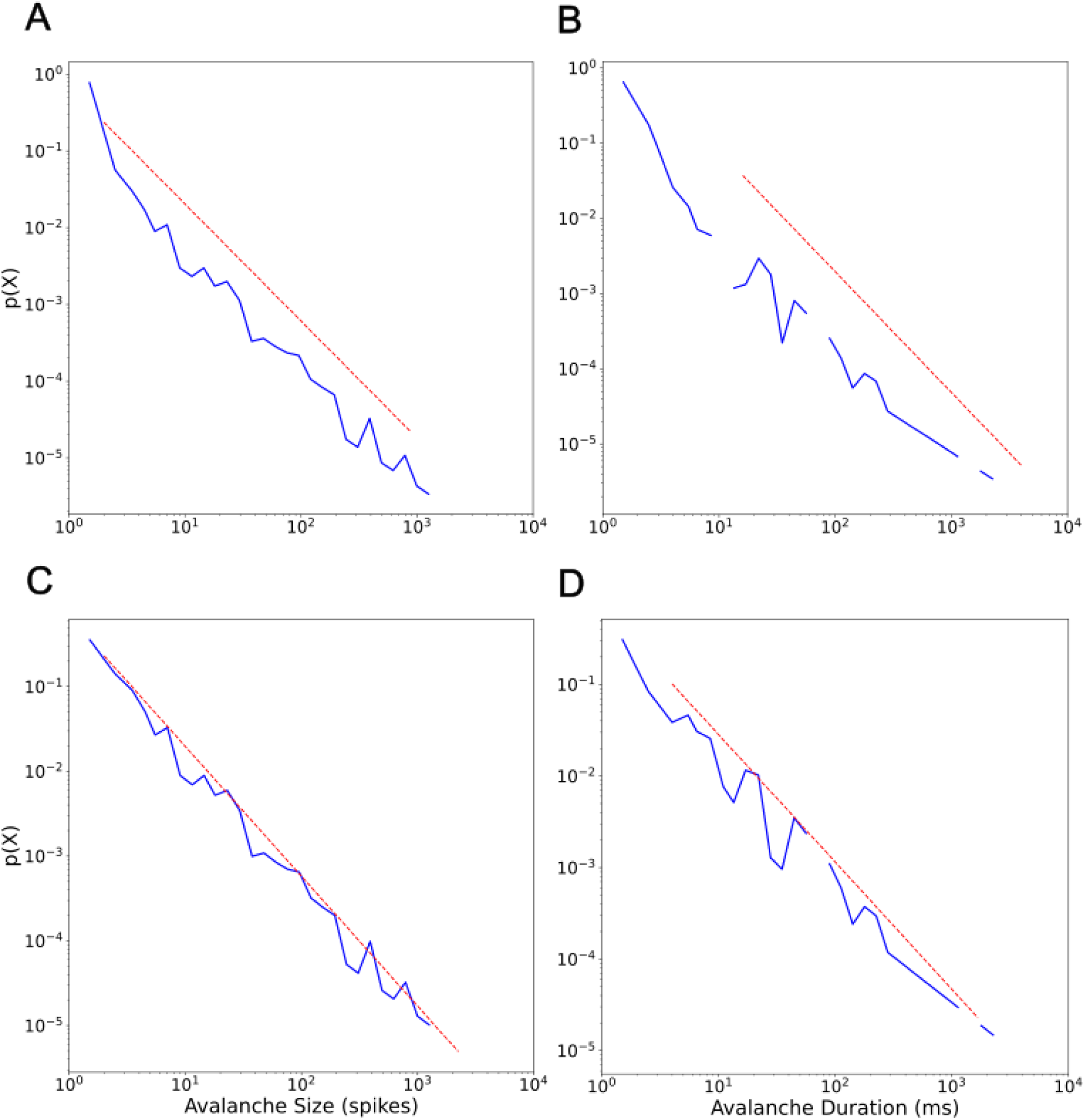
Power-law fits for all neurons in our 10 minute simulation of an M1 cortical column. A. Avalanche size probability density distribution with the number of avalanches normalized (y-axis) and the size of the avalanche (number of spikes; x-axis). Power-law fit equals −1.52 (red dashed line; sigma = 0.035, D = 0.028). B. Avalanche duration probability density distribution showing the number of avalanches normalized (y-axis) and their durations (milliseconds; x-axis). Power-law fit equals −1.61 (red dashed line; sigma = 0.088, D = 0.067). C. Same as A except only Irregular and Fragment avalanches. Power-law fit equals −1.53 (sigma = 0.036, D = 0.033). D. Same as A except only Irregular and Fragment avalanches. Power-law fit equals −1.39 (sigma = 0.045, D = 0.063). Analyzed using the Python powerlaw package ^44^.

### Inter-avalanche intervals

Avalanches are composed of at least 1 spike every millisecond bounded by quiescent periods of 1 or more milliseconds. These quiescent periods between avalanches, or inter-avalanche intervals, typically lasted 1-4 ms but ranged up to about 40 ms (Figs S5, S6, S9, S12, S15, S18, S19). Investigating the quiescent periods revealed sustained depolarization of the apical dendrites, mediated by NMDA receptors, maintained for up to around 60 ms.

### Avalanche types

Every avalanche we observed in this study was composed of one or more of four patterns: irregular, delta, beta, and gamma. The fifteen avalanches in Fig 4A provide a representative sample of the self-organized patterns and pattern composition of avalanches observed in simulated M1. The first row in Fig 4A begins with a short duration, 3 ms, avalanche displaying the irregular pattern with IT5B/IT6 spikes and no inhibitory neuron activity. We have named these Irregular avalanches. Irregular avalanches were relatively small (size range 1 to 109 spikes; Table I) with short durations (duration range 1 to 40 ms; Table I) and composed 65.3% of the total number of avalanches but only 6.0% of total cortical activity duration (Table I). A log-log plot of the Irregular avalanches (Fig 6; light-blue) provides a visualization of their range. The second avalanche in Fig 4A row one is relatively brief, 171 ms, with prominent activity in IT5B/IT6 but also inhibitory activity primarily in SOM2/3 and PV6. We named these Fragment avalanches because they appear mostly broken off of the beginning or end of Delta+ avalanches (defined below). Fragment avalanches compose 30.3% of the total avalanche numbers and 12.0% of total cortical activity duration and range in size to more than 20 times larger than Irregular avalanches and more than 6 times longer duration (Table I; Fig 6, orange). The third avalanche in Fig 4A row one is composed of the delta pattern and a nested gamma pattern. Delta with nested gamma was a commonly observed pattern in M1, which repeated approximately once every second (Fig 1) and was often preceded with beta pattern activity (see beta patterns preceding delta patterns in Fig 4A rows two and three). We named avalanches including the delta pattern Delta Plus (Delta+) avalanches since these avalanches almost always included other patterns in addition to the delta pattern. Delta+ avalanches were distinct in their range of sizes from Irregular and Fragment avalanches, composed just 3.3% of the total avalanche numbers, but were the most commonly active and constituted 79.5% of the total duration of avalanche activity in our representative simulated M1 (Table I; Fig 6, purple). The third avalanche in Fig 4A row one is followed by three Fragment avalanches. The first is clearly broken off from the end of the delta pattern envelope. Fragment avalanches are distinguished from Irregular avalanches by their inclusion of activity from neurons other than IT5B/IT6, including from inhibitory neurons.

**Figure 6.**
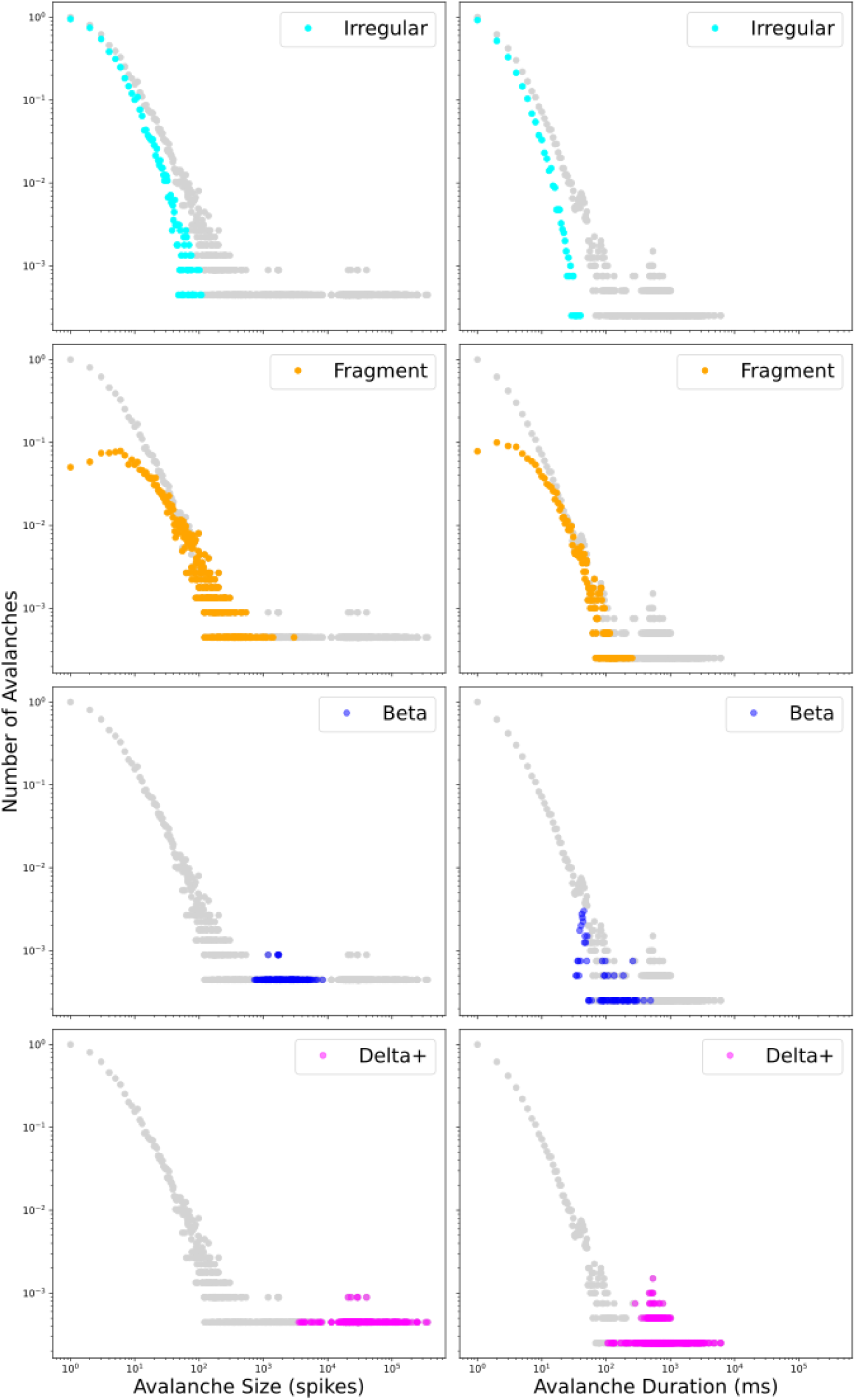
Log-log probability density distributions of avalanche size (first column) and duration (second column) for all avalanches (gray) and each of four avalanche types.

A two period Delta+ avalanche fills all of Fig 4A row two. This avalanche started with beta pattern activity and was followed by delta and nested gamma patterns. The avalanche in row two ended mid-beta activity and was followed by smaller avalanches in row three beginning with a 51 ms avalanche with a single period of beta pattern activity. Avalanches composed entirely of beta patterns are Beta avalanches. Beta avalanches, like the first and fourth avalanches in row three, form their own cluster (Fig 6; blue) that does not overlap with Irregular avalanche sizes but does slightly overlap in duration (Table I). Beta avalanches compose 1.1% of the total avalanche numbers and 2.5% of total duration (Table I). The beta pattern continued in the second avalanche in Fig 4A row three, until the delta pattern appeared with a nested gamma pattern making this second avalanche in row three a Delta+ avalanche.

In total, four types of avalanches were observed. Two types were composed of just one pattern. Irregular avalanches composed of irregular patterns and Beta avalanches composed of beta patterns. One type, the Delta+ avalanche, included all avalanches with any combination of delta and gamma patterns, and sometimes also included any combination of the irregular and beta patterns. And one type, the Fragment avalanche, was composed of fragments of all but Irregular avalanches.

### Avalanche types and power-law distributions

The Irregular and Fragment avalanche stand out in the log-log graphs in Fig 6 as closest in form to a power-law. The power-law fit of the Irregular and Fragment avalanche sizes together is −1.53 (Fig 5C). The power-law fit to all four is −1.52 (Fig 5A). The combination of Irregular and Fragment avalanches gives a power-law value identical or very close to the value obtained from the entire set of avalanches. Beta and Delta+ avalanches form small clusters (Table 1; 163 or 1.1% and 519 or 3.3% of total population of avalanches respectively) in the tail of the power-law graph (Fig 6; blue and purple respectively). Avalanches that include dominant frequency patterns, the Beta and Delta+ avalanches, do not appreciably contribute to the power-law value. It is especially striking that Delta+ avalanches, compose just 3.3% of the total avalanche population but compose the largest chunk of cortical activity 79.5%, and yet its cluster is farthest out on the power-law tail and contributes little, if at all, to the power-law value.

**Table I.** Data from a representative 10 minutes simulation of M1 (sM1_06-15-2019_02).

| Avalanche Type | Number | Percent Total Num | Size Range (spikes) | Size Range (neurons) | Duration Range (ms) | Total Duration (ms) | Percent Total Dur |
| --- | --- | --- | --- | --- | --- | --- | --- |
| Irregular | 10,173 | 65.3% | 1-109 | 1-102 | 1-40 | 34,446 | 6.0% |
| Fragment | 4,723 | 30.3% | 1-2,999 | 1-1,199 | 1-253 | 69,372 | 12.0% |
| Beta | 163 | 1.1% | 728 - 8,346 | 647 - 4,370 | 34 - 493 | 14,607 | 2.5% |
| Delta+ | 519 | 3.3% | 3,536 - 358,184 | 1,132 - 7,599 | 106 - 6,138 | 458,453 | 79.5% |
| Total | 15,578 | 100.0% |  |  |  | 576,878 | 100% |

## Discussion

Size and duration power-law distributions of clustered spiking activity known as avalanches have been widely observed in the cortex ^1,10,11^. In this study, simulated motor cortex frequently displayed robust avalanche activity with power-law values similar to those established by in vitro ^4–7^ and in vivo ^8–18^ experiments. We set out to observe what the well-established phenomenon of an avalanche looked like in its entirety within a volume of cortex.

The data-driven detailed simulations of motor cortex displayed consistent and highly-structured patterns of self-organized and self-sustained activity to a brief (100 ms, 0.57 nA/cell) and unstructured stimulation to less than 2% of the more than 10,000 neurons. We observed four patterns of activity organized across the simulated motor cortex: irregular and the three rhythm dominated patterns delta, beta, and gamma. Once self-organized and self-sustained activity appeared in our simulated slice there was always, at minimum, irregular activity composed of one, the other, or both layer 5B and layer 6 (IT5B/IT6) excitatory pyramidal neurons (Fig 1). More often, cortical activity included bursts of spikes that traveled across layers and different neuron types. Depending on the pattern of signal flow, this form of activity resulted in the appearance of one or more of the rhythm dominated patterns delta, beta, and gamma (Fig 4). Each of the rhythm dominated patterns included both excitatory and inhibitory neurons.

At the macroscopic level we observed prominent bursts of activity about once every second (1 Hz) or delta activity (Fig 1). A recent study of spontaneous cortical activity using voltage-sensitive dye observed about one spontaneous event every second (1 Hz) in both awake and anesthetized mice ^45^. Similar macroscopic activity patterns were observed by Markram et al ^46^ while investigating a detailed data-driven model of mouse cortex. They showed that slow oscillatory bursting appeared when extracellular calcium concentration was high in the physiological range and, conversely, neuronal activity became asynchronous and irregular at relatively low extracellular calcium concentrations.

Each avalanche was composed of one or more of the 4 cross-cortical spiking patterns and belonged to one of 4 distinct clusters in log-log avalanche probability density distribution plots for size and duration (Fig 6). The 4 avalanche types, one for each cluster, were Irregular, Fragment, Beta, and Delta+. Irregular avalanches were composed of the irregular spiking pattern from one or both IT5B/IT6 excitatory pyramidal neurons. Fragment avalanches were often composed of a broken off fragment of rhythm dominated avalanches and, unlike Irregular avalanches, included inhibitory neuron activity. Beta avalanches contained one or more periods of beta spiking patterns. Delta+ avalanches included all avalanches with delta patterns, many of which included nested gamma spiking patterns, and some of which included irregular and/or beta spiking patterns.

Interestingly, in vivo local field potential (LFP) data from cat and monkey primary visual cortex had been analyzed using principal component analysis (PCA) and the first three principal components (PC) composed each segment of a three dimensional PC space that displayed a characteristic pattern of segment clusters across cat and monkey datasets ^20^. Using k-means cluster analysis the research team extracted different clusters of similar frequency compositions with the optimal number of distinct clusters around 5 for cat and 4 for monkey. We see 4 distinct clusters of avalanche types in our M1 simulation data with Delta+ avalanches active most of the time (79.5% for our representative case, Table I). Similarly, cortical activity was active with one cluster type 58% of the time in cats and 75% of the time in monkeys with the other cortical dynamics (represented by the other cluster types) active much less frequently.

Neurophysiologists have begun to approach signals in the brain not readily attributable to particular sensory or motor events as self-generated and self-organized signals crucial to brain function rather than noise to be ignored or averaged out. Ongoing cortical activity in the absence of stimuli has been observed to be comparable with stimulus driven activity and has been correlated with behavior ^37,47–52^. We demonstrate that so-called spontaneous activity in a data-driven detailed cortical simulation is not random but is organized into structured spatio-temporal profiles that reflect the functional architecture of the cortex.

In conclusion, we have viewed four self-organized and self-sustained spiking patterns in the simulated motor cortex and their organization into four classes of avalanches. A majority of avalanches are Irregular avalanches (65.3%) composed mostly of deep excitatory pyramidal neurons from layers 5B and 6 (IT5B/IT6). The combination of Irregular and Fragment avalanches (Fragment: 30.3% of the total number of avalanches; Irregular + Fragment: 95.6%) are dominant in determining the power-law value for avalanche size and duration. However, even though together Irregular and Fragment avalanches were 95.6% of the total number of avalanches, they made up only 18.0% of the total duration of avalanche, and therefore cortical, activity. The two observed rhythm dominated avalanches, Beta and Delta+, composed only 4.4% of the total number of avalanches but made up 82.0% of the total duration of avalanche activity. The fact that cortical activity in our data-driven simulation of motor cortex expressed dominant rhythmic activity most of the time (82%), driven by Beta and Delta+ avalanches which composed only 4.4% of avalanches, presents interesting questions related to how the brain may process signals in self-sustained and self-organized spontaneous activity similar to our observations. We hypothesize that the nonrhythmic Irregular and Fragment avalanches provide the decorrelated spike fluctuations that result in increased entropy and reduced redundancy that are thought to maximize information processing and storage capacity ^12,28,29^. While the rhythm dominated Beta and Delta+ avalanches may provide temporal coordination across neurons, time segmentation for information transfer, and signal amplification through frequency resonance ^53–55^.

## Acknowledgments

We gratefully acknowledge discussions with Dietmar Plenz.

This research was funded by the National Institute of Neurological Disorders and Stroke (grant#: R01 EB022903, U01 EB017695). This work used Stampede2 at Texas Advanced Computing Center through allocation TG-IBN160014 from XSEDE, which is now the Advanced Cyberinfrastructure Coordination Ecosystem: Services & Support (ACCESS) program, supported by National Science Foundation grants #2138259, #2138286, #2138307, #2137603, and #2138296. For the purpose of open access, the author has applied a CC BY public copyright license to all Author Accepted Manuscripts arising from this submission.

## Author Contributions

D.W.D. and W.W.L. contributed to the conception and design of the study; S.D-B contributed to the conception of the study; D.W.D. contributed to the acquisition and data analysis; J.J. contributed to data analysis. D.W.D. and W.W.L. contributed to preparing the figures; D.W.D. and W.W.L. contributed to drafting the text.

## Potential Conflicts of Interest

Nothing to report.

## Materials and Methods

The methods below describe our data-driven model development and key features of the final model.

### Morphology and physiology of neuron classes

Seven excitatory pyramidal cells and two interneuron cell models were employed in the network. In previous work we developed layer 5B PT corticospinal cell and L5 IT corticostriatal cell models that reproduced in vitro electrophysiological responses to somatic current injections, including sub- and super-threshold voltage trajectories and f-I curves ^54,56^. To achieve this, we optimized the parameters of the Hodgkin-Huxley neuron model ionic channels – Na, Kdr, Ka, Kd, HCN, CaL, CaN, KCa – within a range of values constrained by the literature. The corticospinal and corticostriatal cell model morphologies had 706 and 325 compartments, respectively, digitally reconstructed from 3D microscopy images. Morphologies are available via NeuroMorpho.org ^57^; archive name “Suter Shepherd”). For the current simulations, we further improved the PT model by 1. increasing the concentration of Ca2+ channels (“hot zones”) between the nexus and apical tuft, following parameters published in ^58^; 2. lowering dendritic Na+ channel density in order to increase the threshold required to elicit dendritic spikes, which then required adapting the axon sodium conductance and axial resistance to maintain a similar f-I curve; 3. replacing the HCN channel model and distribution with a more recent implementation ^59^.The new HCN channel reproduced a wider range of experimental observations than our previous implementation ^60^, including the change from excitatory to inhibitory effect in response to synaptic inputs of increasing strength ^61^. This was achieved by including a shunting current proportional to Ih. We tuned the HCN parameters (lk and vrevlk) and passive parameters to reproduce the findings noted above, while keeping a consistent f-I curve ^56^.

The network model includes five other excitatory cell classes: layer 2/3, layer 4, layer 5B and layer 6 IT neurons and layer 6 CT neurons. These cell classes were implemented using simpler models as a trade-off to enable running a larger number of exploratory network simulations. Previously we had optimized 6-compartment neuron models to reproduce somatic current clamp recordings from two IT cells in layers 5A and 5B. The layer 5A cell had a lower f-I slope (77 Hz/nA) and higher rheobase (250 nA) than that in layer 5B (98 Hz/nA and 100 nA). Two broad IT categories based on projection and intrinsic properties: corticocortical IT cells found in upper layers 2/3 and 4 which exhibited a lower f-I slope (~72 Hz/nA) and higher rheobase (~281 pA) than IT corticostriatal cells in deeper layers 5A, 5B and 6 (~96 Hz/nA and ~106 pA) ^56,62,63^. CT neurons’ f-I rheobase and slope (69 Hz/nA and 298 pA) was closer to that of corticocortical neurons ^63^. We therefore employed the layer 5A IT model for layers 2/3 and 4 IT neurons and layer 6 CT neurons, and the layer 5B IT model for layers 5A, 5B and 6 IT neurons. We further adapted cell models by modifying their apical dendrite length to match the average cortical depth of the layer, thus introducing small variations in the firing responses of neurons across layers.

We implemented models for two major classes of GABAergic interneurons ^64^: parvalbumin-expressing fast-spiking (PV) and somatostatin-expressing low-threshold spiking neurons (SOM). We employed existing simplified 3-compartment (soma, axon, dendrite) models ^65^ and increased their dendritic length to better match the average f-I slope and rheobase experimental values of cortical basket (PV) and Martinotti (SOM) cells (Neuroelectro online database; ^66^.

### Microcircuit composition: neuron locations, densities and ratios

We modeled a cylindrical volume of the mouse M1 cortical microcircuit with a 300 μm diameter and 1350 μm height (cortical depth) at full neuronal density for a total of 10,073 neurons. Cylinder diameter was chosen to approximately match the horizontal dendritic span of a corticospinal neuron located at the center, consistent with the approach used in the Human Brain Project model of the rat S1 microcircuit ^46^. Mouse cortical depth and boundaries for layers 2/3, 4, 5A, 5B and 6 were based on published experimental data ^62,67,68^. Although traditionally M1 has been considered an agranular area lacking layer 4, recently identified M1 pyramidal neurons with the expected prototypical physiological, morphological and wiring properties of layer 4 neurons ^62,69,70^, and therefore incorporated this layer in the model.

Cell classes present in each layer were determined based on mouse M1 studies ^56,62–65,68,71^. IT cell populations were present in all layers, whereas the PT cell population was confined to layer 5B, and the CT cell population only occupied layer 6. SOM and PV interneuron populations were distributed in each layer. Neuronal densities (neurons per mm3) for each layer were taken from a histological and imaging study of mouse agranular cortex ^72^. The proportion of excitatory to inhibitory neurons per layer was obtained from mouse S1 data ^73^. The proportion of IT to PT and IT to CT cells in layers 5B and 6, respectively, were both estimated as 1:1 ^56,64,74^. The ratio of PV to SOM neurons per layer was estimated as 2:1 based on mouse M1 and S1 studies ^75,76^. Since data for M1 layer 4 was not available, interneuron populations labeled PV5A and SOM5A occupy both layers 4 and 5A. The number of cells for each population was calculated based on the modeled cylinder dimensions, layer boundaries and neuronal proportions and densities per layer.

### Local connectivity

We calculated local connectivity between M1 neurons by combining data from multiple studies. Data on excitatory inputs to excitatory neurons (IT, PT and CT) was primarily derived from mapping studies using whole-cell recording, glutamate uncaging-based laser-scanning photostimulation (LSPS) and subcellular channelrhodopsin-2-assisted circuit mapping (sCRACM) analysis ^62,67,68,74^. Connectivity data was postsynaptic cell class-specific and employed normalized cortical depth (NCD) instead of layers as the primary reference system. Unlike layer definitions which can be interpreted differently between studies, NCD provides a well-defined, consistent and continuous reference system, depending only on two readily-identifiable landmarks: pia (NCD=0) and white matter (NCD=1). Incorporating NCD-based connectivity into our model allowed us to capture wiring patterns down to a 100 μm spatial resolution, well beyond traditional layer-based cortical models. M1 connectivity varied systematically within layers. For example, the strength of inputs from layer 2/3 to L5B corticospinal cells depends significantly on cell soma depth, with upper neurons receiving much stronger input ^68^.

Connection strength thus depended on presynaptic NCD and postsynaptic NCD and cell class. For postsynaptic IT neurons with NCD ranging from 0.1 to 0.37 (layers 2/3 and 4) and 0.8 to 1.0 (layer 6) we determined connection strengths based on data from ^67^ with cortical depth resolution of 140 μm-resolution. For postsynaptic IT and PT neurons with NCD between 0.37 and 0.8 (layers 5A and 5B) we employed connectivity strength data from ^68^ with cortical depth resolution of 100 μm. For postsynaptic CT neurons in layer 6 we used the same connection strengths as for layer 6 IT cells ^67^, but reduced to 62% of original values, following published data on the circuitry of M1 CT neurons ^74^. Data ^74^ also suggested that connection strength from layer 4 to layer 2/3 IT cells was similar to that measured in S1, so for these projections we employed values from Lefort’s S1 connectivity strength matrix ^73^. Experimentally, these connections were found to be four times stronger than in the opposite direction – from layer 2/3 to layer 4 – so we decreased the latter in the model to match this ratio.

Following previous publications ^73,77^ we defined connection strength (scon, in mV) between two populations, as the product of their probability of connection (pcon) and the unitary connection somatic EPSP amplitude in mV (vcon), i.e. scon = pcon × vcon. We employed this equivalence to disentangle the connection scon values provided by the above LSPS studies into pcon and vcon values that we could use to implement the model. First, we rescaled the LSPS raw current values in pA ^62,67,68,74^ to match scon data from a paired recording study of mouse M1 L5 excitatory circuits ^77^. Next, we calculated the M1 NCD-based vcon matrix by interpolating a layerwise unitary connection EPSP amplitude matrix of mouse S1 ^73^, and thresholding values between 0.3 and 1.0 mV. Finally, we calculated the probability connection matrix as pcon = scon/vcon.

To implement vcon values in the model we calculated the required NEURON connection weight of an excitatory synaptic input to generate a somatic EPSP of 0.5 mV at each neuron segment. This allowed us to calculate a scaling factor for each segment that converted vcon values into NEURON weights, such that the somatic EPSP response to a unitary connection input was independent of synaptic location. This is consistent with experimental evidence showing synaptic conductances increased with distance from soma, to normalize somatic EPSP amplitude of inputs within 300 μm of soma ^78^. Following this study, scaling factor values above 4.0 – such as those calculated for PT cell apical tufts – were thresholded to avoid overexcitability in the network context where each cell receives hundreds of inputs that interact nonlinearly ^79,80^. For morphologically detailed cells (layer 5A IT and layer 5B PT), the number of synaptic contacts per unitary connection (or simply, synapses per connection) was set to five, an estimated average consistent with the limited mouse M1 data ^81^ and rat S1 studies ^46,82^. Individual synaptic weights were calculated by dividing the unitary connection weight (vcon) by the number of synapses per connection. Although the method does not account for nonlinear summation effects ^80^, it provides a reasonable approximation and enables employing a more realistic number and spatial distribution of synapses, which may be key for dendritic computations ^83^. For the remaining cell models, all with six compartments or less, a single synapse per connection was used.

For excitatory inputs to inhibitory cell types (PV and SOM) we started with the same values as for IT cell types but adapted these based on the specific connectivity patterns reported for mouse M1 interneurons ^74,84^. Following the layer-based description in these studies, we employed three major subdivisions: layer 2/3 (NCD 0.12 to 0.31), layers 4, 5A and 5B (NCD 0.31 to 0.77) and layer 6 (NCD 0.77 to 1.0). We increased the probability of layer 2/3 excitatory connections to layers 4, 5A and 5B SOM cells by 50% and decreased that to PV cells by 50% ^84^. We implemented the opposite pattern for excitatory connections arising from layer 4, 5A, 5B IT cells such that PV interneurons received stronger intralaminar inputs than SOM cells ^84^. The model also accounts for layer 6 CT neurons generating relatively more inhibition than IT neurons ^74^. Inhibitory connections from interneurons (PV and SOM) to other cell types were limited to neurons in the same layer ^76^, with layers 4, 5A, and 5B combined into a single layer ^71^. Probability of connection decayed exponentially with the distance between the pre- and post-synaptic cell bodies with length constant of 100 μm ^85,86^. We introduced a correction factor to the distance-dependent connectivity measures to avoid the border effect, i.e., cells near the modeled volume edges receiving less or weaker connections than those in the center.

For comparison with other models and experiments, we calculated the probability of connection matrices arranged by population (instead of NCD) for the base model network instantiation used throughout the results.

Excitatory synapses consisted of colocalized AMPA (rise, decay τ: 0.05, 5.3 ms) and NMDA (rise, decay τ: 15, 150 ms) receptors, both with reversal potential of 0 mV. The ratio of NMDA to AMPA receptors was 1.0 ^87^, meaning their weights were each set to 50% of the connection weight. NMDA conductance was scaled by 1/(1 + 0.28 · M g · exp (−0.062 · V)); Mg = 1mM 61. Inhibitory synapses from SOM to excitatory neurons consisted of a slow GABAA receptor (rise, decay τ: 2, 100 ms) and GABAB receptor, in a 90% to 10% proportion; synapses from SOM to inhibitory neurons only included the slow GABAA receptor; and synapses from PV to other neurons consisted of a fast GABAA receptor (rise, decay τ: 0.07, 18.2). The reversal potential was −80 mV for GABAA and −95 mV for GABAB. The GABAB synapse was modeled using second messenger connectivity to a G protein-coupled inwardly-rectifying potassium channel (GIRK) ^88^. The remaining synapses were modeled with a double-exponential mechanism.

Connection delays were estimated as 2 ms plus a variable delay depending on the distance between the pre- and postsynaptic cell bodies assuming a propagation speed of 0.5 m/s.

### Dendritic distribution of synaptic inputs

Experimental evidence demonstrates the location of synapses along dendritic trees follows very specific patterns of organization that depend on the brain region, cell type and cortical depth ^89,90^; these are likely to result in important functional effects ^80,91,92^. Synaptic locations were automatically calculated for each cell based on its morphology and the pre- and postsynaptic cell type-specific radial synaptic density function. Synaptic inputs from PV to excitatory cells were located perisomatically (50 μm around soma); SOM inputs targeted apical dendrites of excitatory neurons ^71,76^; and all inputs to PV and SOM cells targeted apical dendrites. For projections where no data synaptic distribution data was available – IT/CT to IT/CT cells – we assumed a uniform dendritic length distribution.

### Model implementation, simulation and analysis

The model was developed using parallel NEURON (neuron.yale.edu) ^93^ and NetPyNE (www.netpyne.org) ^94^, a Python package to facilitate the development of biological neuronal networks in the NEURON simulator. NetPyNE emphasizes the incorporation of multiscale anatomical and physiological data at varying levels of detail. It converts a set of simple, standardized high-level specifications in a declarative format into a NEURON model. This high-level language enables, for example, defining connectivity as a function of NCD, and distributing synapses across neurons based on normalized synaptic density maps. NetPyNE facilitates running parallel simulations by taking care of distributing the workload and gathering data across computing nodes, and automates the submission of batches of simulations for parameter optimization and exploration. It also provides a powerful set of analysis methods so the user can plot spike raster plots, LFP power spectra, information transfer measures, connectivity matrices, or intrinsic time-varying variables (eg. voltage) of any subset of cells. To facilitate data sharing, the package saves and loads the specifications, network, and simulation results using common file formats (Pickle, Matlab, JSON or HDF5), and can convert to and from NeuroML ^95,96^ and SONATA ^97^, standard data formats for exchanging models in computational neuroscience. Simulations were run on XSEDE supercomputers Comet and Stampede 2 and on Google Cloud supercomputers.

### Cortical column configuration

The M1 cortical column model was 300 um in diameter and 1,350 um deep (from pia to white matter). The volume of the cortical column was about 95,425,877 um^3^. Stimulating electrodes applied current to a 40 um diameter area tangential to the cortical column surface and through its entire depth. The volume stimulated by a stimulating electrode is about 1,696,460 um^3^. A stimulating electrode applies current to about 1.8% of the total volume of simulated M1.

The neuronal firing rates across the entire cortical column population gave a lognormal distribution with a power curve distribution of neurons with very low firing rates from about 0.003 Hz to about 0.05 Hz (Fig S1). In vivo experiments may not record from these neurons that spike about once every 20 seconds to about once every 6 minutes and they comprise 239 out of 7,935 spike trains collected or 3 percent of the spiking population of neurons.

### Stimulation and data collection

We briefly stimulated our isolated M1 slice (no afferent or efferent connections) for 100 ms with a 0.57 nA current using a simulated stimulating electrode placed perpendicular to the cortical surface and applying current to any neuron within a 40 *μ*m diameter (centered on the electrode) from the cortical surface to white matter.

Seven cortical columns were constructed using identical wiring densities between neuronal populations but unique connectivity defined using random seeds. In each of the 7 unique cortical columns we tested 49 locations in a 7 by 7 grid of points across the surface of M1, each point separated from the other by 40 *μ*m. A data set consisted of 49 activity records, each recorded for 60 seconds (60,000 ms) from the grid. From these data we selected one stimulus location from each unique cortical column to run a 10 minute simulation. Six of these showed full cross-layer activity and were therefore used for further analysis (1 column showed in only IT5B/IT6 and was excluded).

### Power-law analysis

We analyzed our avalanche data for power-law distributions using the Python powerlaw package ^44^.

### SPUD analysis

We extracted the three dominant frequency patterns using a time-domain features algorithm known as SPUD ^43^. Delta waves were selected from large IT2/3 activity peaks (>= 250 spikes) from 1ms binned histograms of cortical column activity. IT2/3 peaks less than or equal to 50ms apart were filtered out. Beta waves were selected using SPUD analysis by picking IT5B peak activity (>= 15 spikes) from cortical activity and filtering IT5B activity that occurred during robust IT2/3 activity (>= 250 spikes). Gamma waves were selected by picking PT5B peak activity (>= 20 spikes) from cortical activity and filtering IT2/3 activity that occurred during robust PT5B activity (>= 250 spikes).

## Supplementary Data

**Figure S1.**
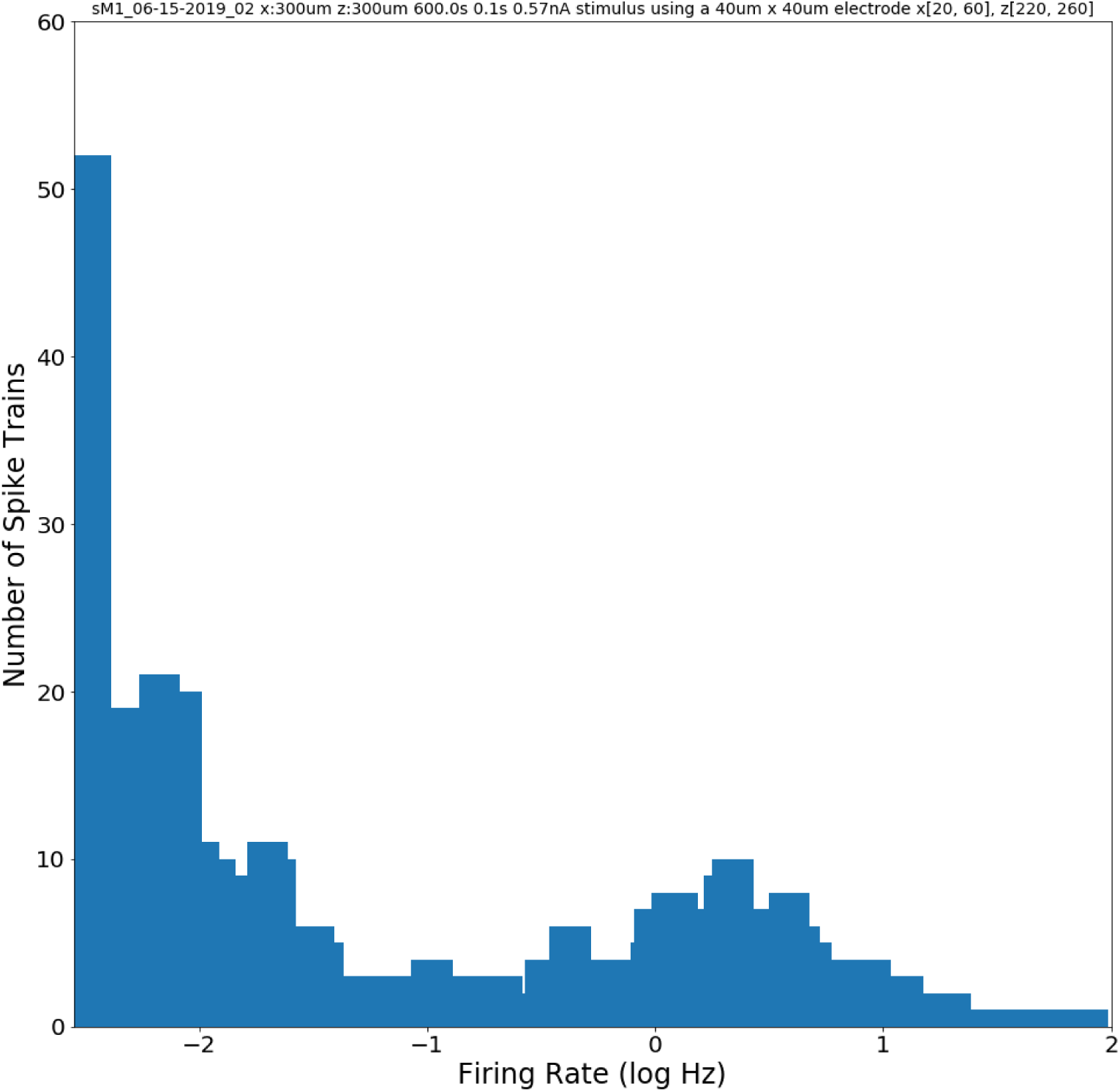
The number of spike trains with logarithm of firing rate (Hertz) for the entire population of neurons. The right hand side shows the lognormal distribution of firing rates typical of firing rate data collected in vivo (from about −0.5 log or 0.32 Hz to about 1.8 log or 63 Hz). At left are very low firing rates (from about −2.5 log or 0.003 Hz to about −1.3 log or 0.05 Hz) that form a power curve distribution. Simulation sM1_06-15-2019_02. Row 2 in Fig S2.

**Figure S2.**
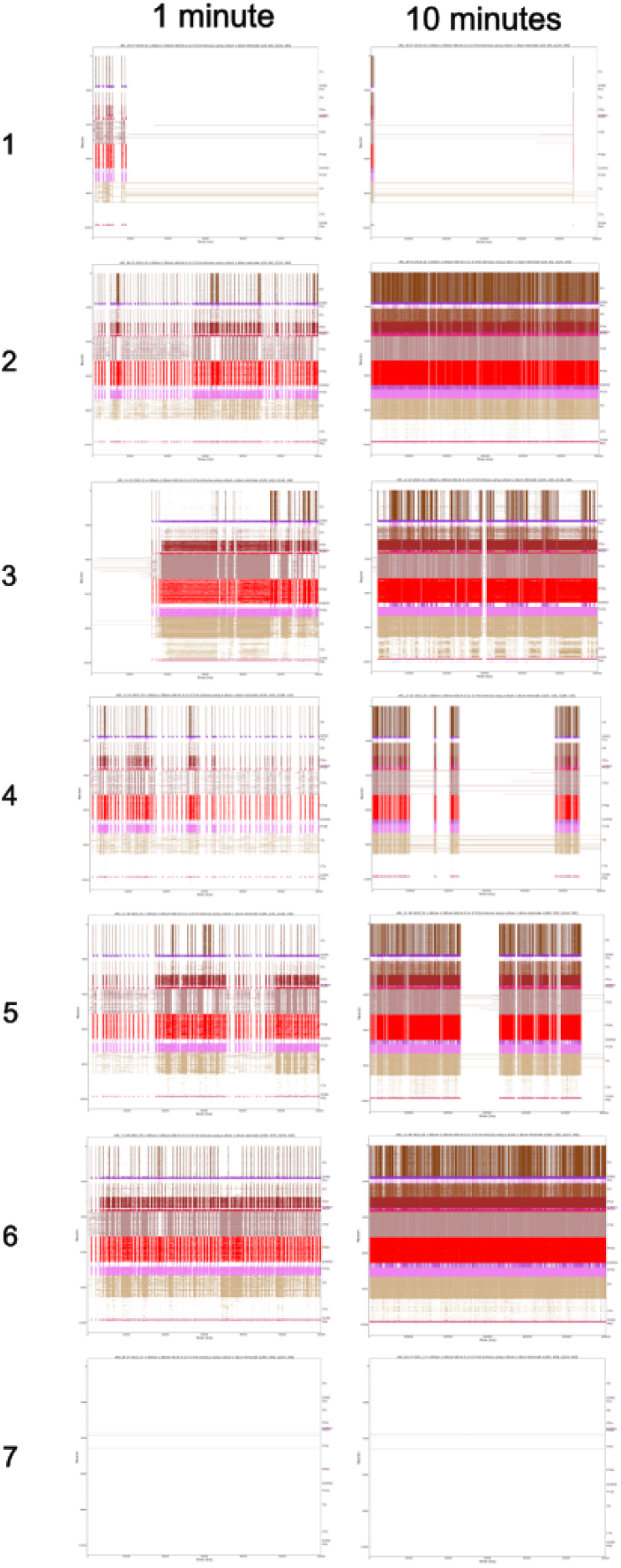
Seven simulations of 10 min cortical activity, each with different connectivities (different connectivity each row). Left column shows raster plots of all neurons for 1 min. Right column shows raster plots of all neurons for 10 min. Row 1: sM1_05-27-2019_01, 2: sM1_06-15-2019_02, 3: sM1_11-22-2020_01, 4: sM1_11-23-2020_05, 5: sM1_11-26-2020_01, 6: sM1_11-28-2020_05, 7: sM1_04-27-2021_17.

**Table S1.** Seven simulations from Fig S2 with values under Row matching the row labels in Fig S2. Simulation: simulation name. Seed: random number seeds provided for ‘conn’ (connections), ‘stim’ (stimulus), and ‘loc’ (locations). Duration (ms): simulation duration in milliseconds. Avalanches: total number of avalanches across the full duration of the simulation.

| Row | Simulation | Seed | Duration (ms) | Avalanches |
| --- | --- | --- | --- | --- |
| 1 | sM1_05-27-2019_01 | cfg.seeds = {'conn': 4321, 'stim': 1234, 'loc': 4321} | 600,000 | 93,020 |
| 2 | sM1_06-15-2019_02 | cfg.seeds = {'conn': 5294, 'stim': 3812, 'loc': 6017} | 600,000 | 15,578 |
| 3 | sM1_11-22-2020_01 | cfg.seeds = {'conn': 3830, 'stim': 3084, 'loc': 3091} | 600,000 | 9,750 |
| 4 | sM1_11-23-2020_05 | cfg.seeds = {'conn': 9364, 'stim': 7334, 'loc': 3329} | 600,000 | 102,739 |
| 5 | sM1_11-26-2020_01 | cfg.seeds = {'conn': 2484, 'stim': 9931, 'loc': 8516} | 600,000 | 38,306 |
| 6 | sM1_11-28-2020_05 | cfg.seeds = {'conn': 4224, 'stim': 4095, 'loc': 7058} | 600,000 | 2,039 |
| 7 | sM1_04-27-2021_17 | cfg.seeds = {'conn': 3209, 'stim': 3639, 'loc': 4626} | 600,000 | 32,969 |

**Table S2.** Data from a 10 minutes simulation of M1 (sM1_05-27-2019_01). Row 1 in Fig S2.

| Avalanche Type | Number | Percent Total Num | Size Range (spikes) | Size Range (neurons) | Duration Range (ms) | Total Duration (ms) | Percent Total Dur |
| --- | --- | --- | --- | --- | --- | --- | --- |
| Irregular | 92,844 | 99.81% | 1-55 | 1-55 | 1-21 | 126,303 | 95.0% |
| Fragment | 166 | 0.18% | 1-434 | 1-281 | 1-94 | 1,165 | 1.0% |
| Beta |  |  |  |  |  |  |  |
| Delta+ | 10 | 0.01% | 5,934-61,194 | 1,585-7,424 | 220-1,372 | 5,364 | 4.0% |
| Total | 93,020 | 100.0% |  |  |  | 132,832 | 100.0% |

**Figure S3.**
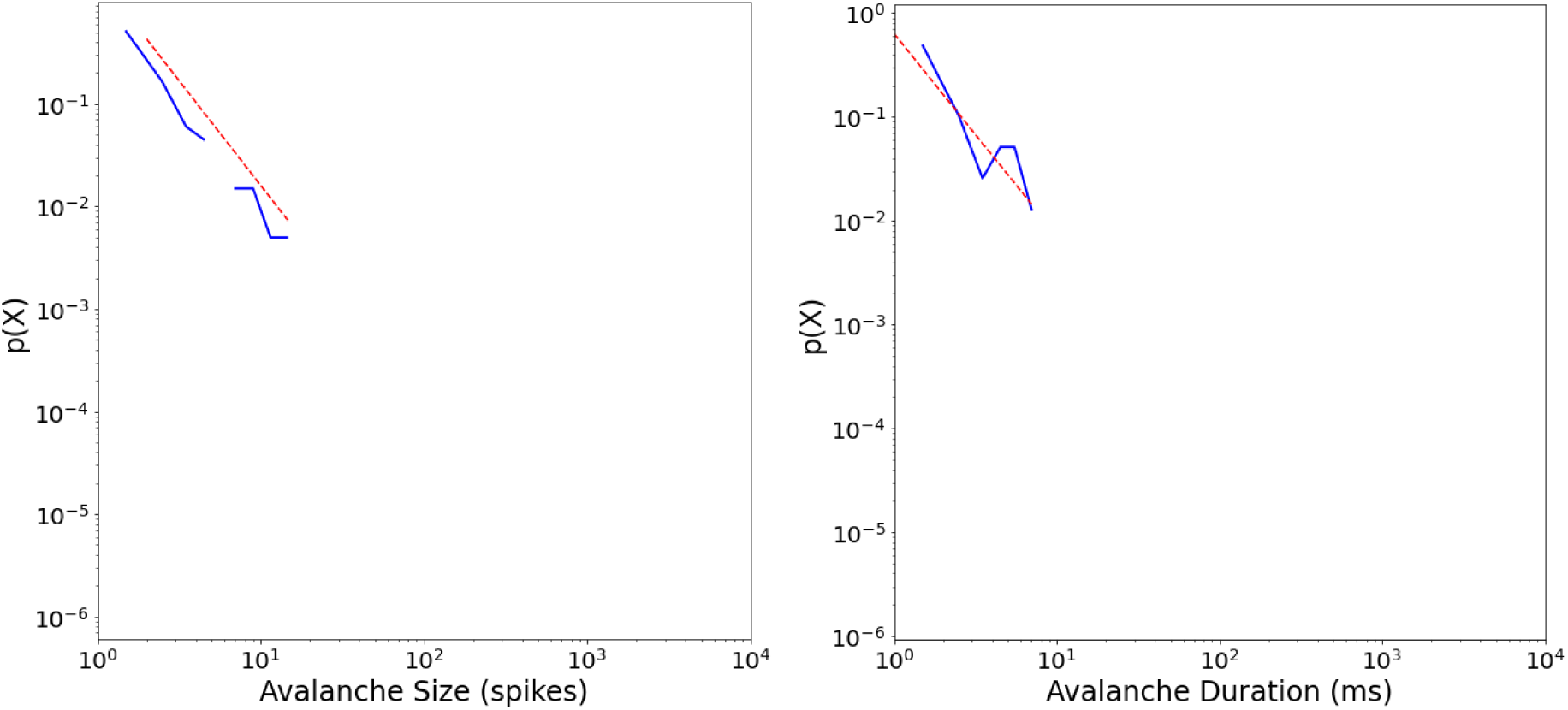
Power-law fits for avalanche size to all neurons in our 10 minute simulation of an M1 cortical column (sM1_05-27-2019_01). Left: Avalanche size probability density distribution with the number of avalanches normalized (y-axis) and the size of the avalanche (number of spikes; x-axis). Power-law fit equals −2.04 (red dashed line; sigma = 0.21, D = 0.085). Right: Avalanche duration probability density distribution showing the number of avalanches normalized (y-axis) and their durations (milliseconds; x-axis). Power-law fit equals −1.94 (red dashed line; sigma = 0.18, D = 0.055). Analyzed using the Python powerlaw package ^44^.

**Figure S4.**
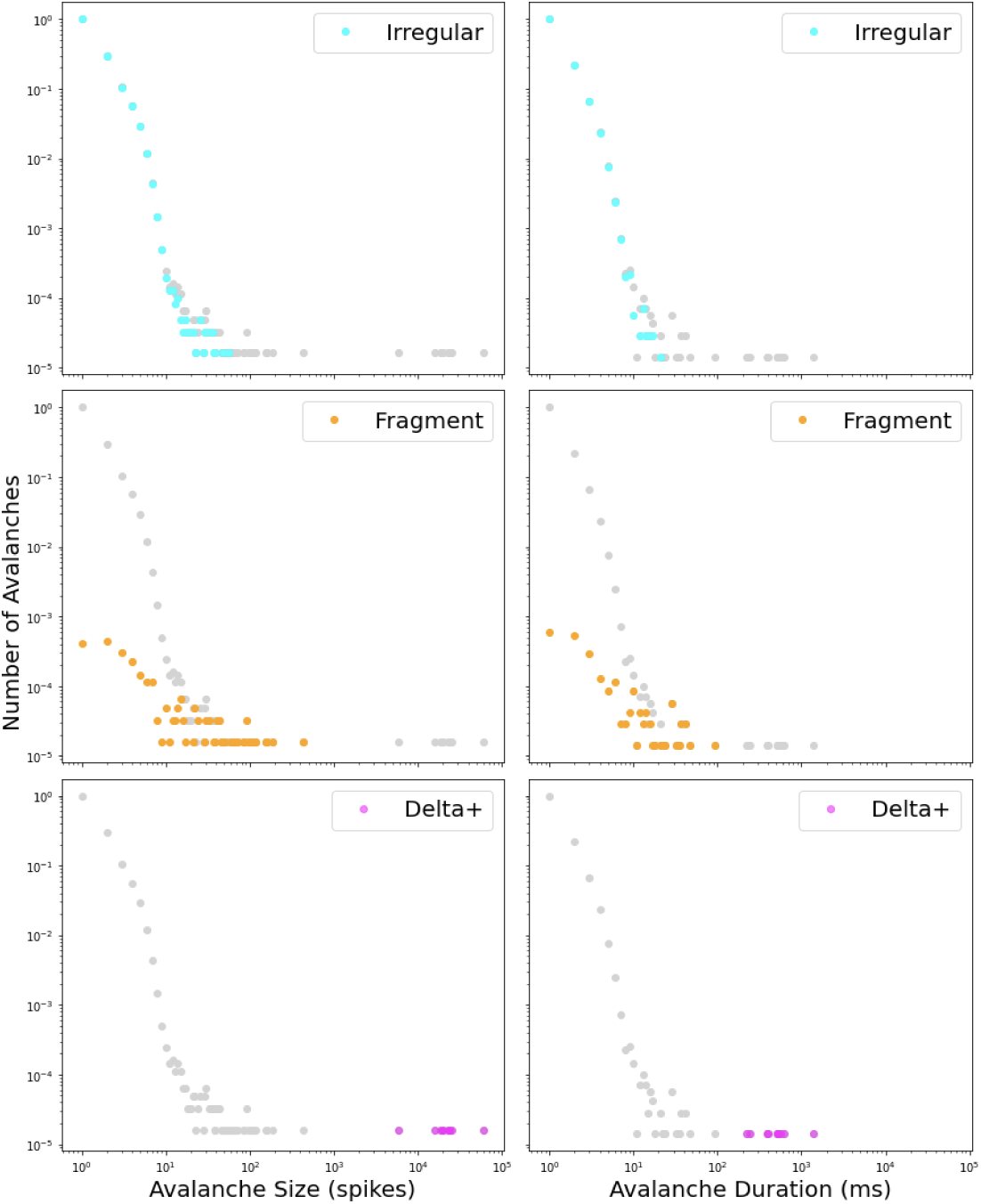
Log-log probability density distributions of avalanche size (first column) and duration (second column) for all avalanches (gray) and each of the three avalanche types seen in this simulation (sM1_05-27-2019_01; row 1; Fig S2).

**Figure S5.**
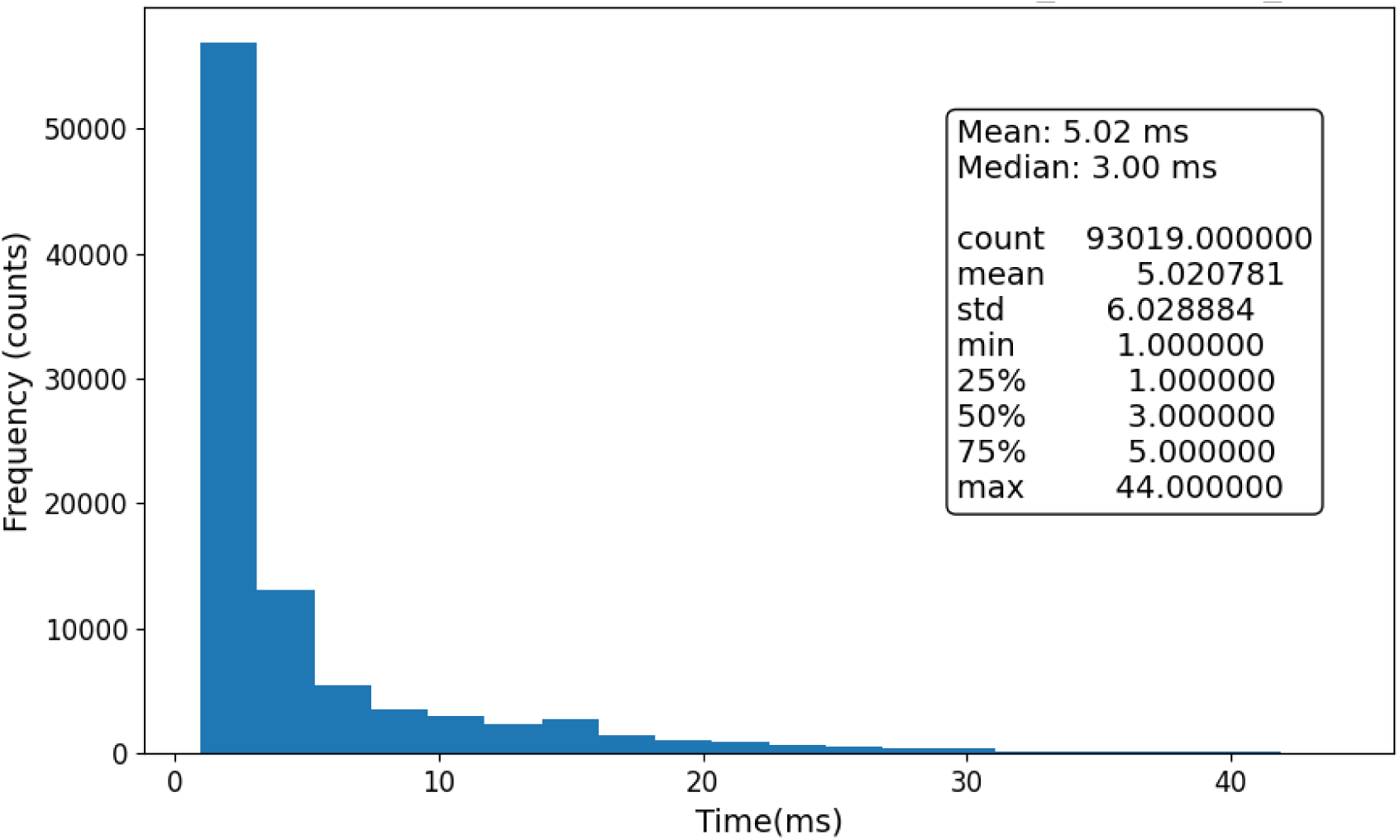
Inter-avalanche intervals in milliseconds (sM1_05-27-2019_01; row 1; Fig S2).

**Figure S6.**
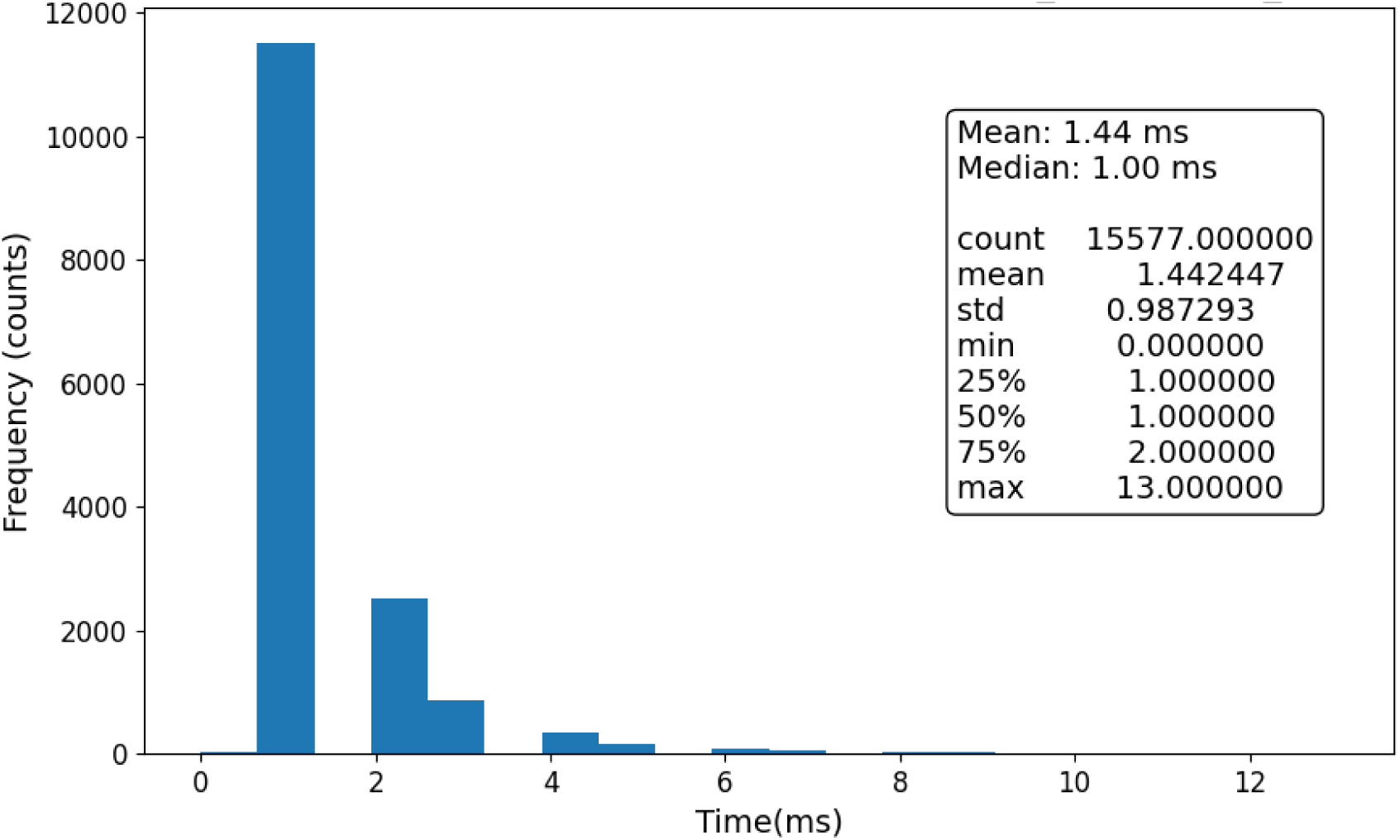
Inter-avalanche intervals in milliseconds (sM1_06-15-2019_02; row 2; Fig S2).

**Table S3.**
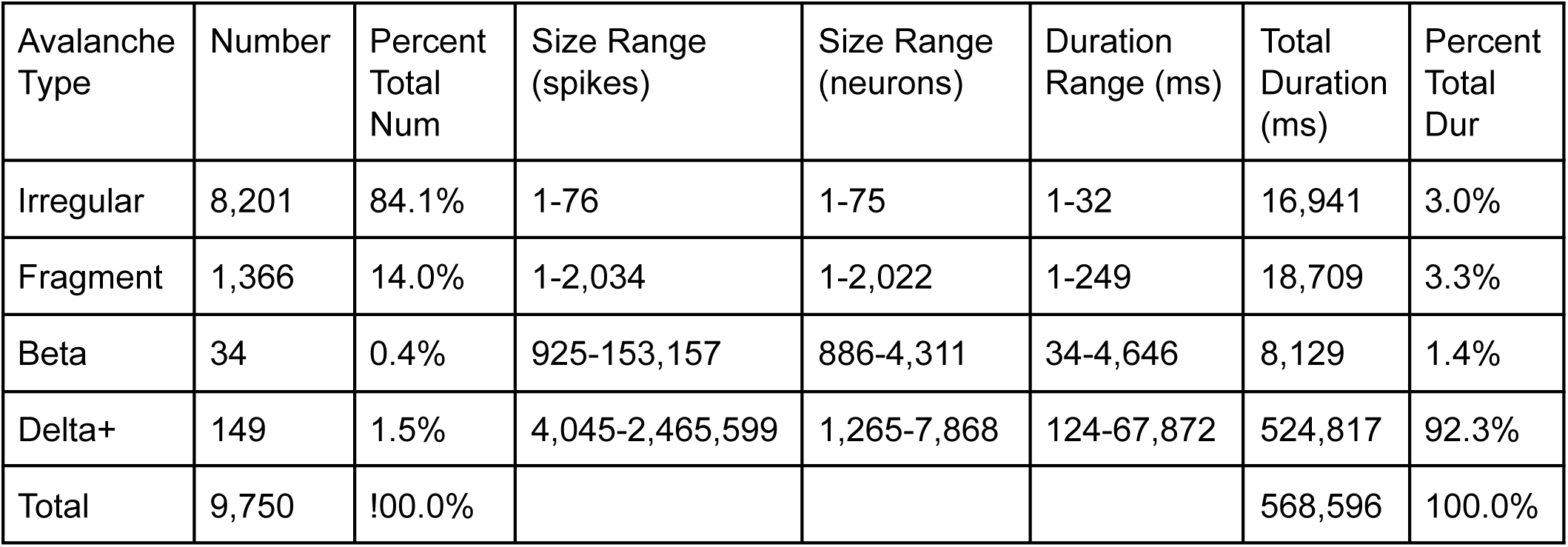
Data from a 10 minutes simulation of M1 (sM1_11–22-2020_01). Row 3 in Fig S2.

| Avalanche Type | Number | Percent Total Num | Size Range (spikes) | Size Range (neurons) | Duration Range (ms) | Total Duration (ms) | Percent Total Dur |
| --- | --- | --- | --- | --- | --- | --- | --- |
| Irregular | 8,201 | 84.1% | 1-76 | 1-75 | 1-32 | 16,941 | 3.0% |
| Fragment | 1,366 | 14.0% | 1-2,034 | 1-2,022 | 1-249 | 18,709 | 3.3% |
| Beta | 34 | 0.4% | 925-153,157 | 886-4,311 | 34-4,646 | 8,129 | 1.4% |
| Delta+ | 149 | 1.5% | 4,045-2,465,599 | 1,265-7,868 | 124-67,872 | 524,817 | 92.3% |
| Total | 9,750 | 100.0% |  |  |  | 568,596 | 100.0% |

**Figure S7.**
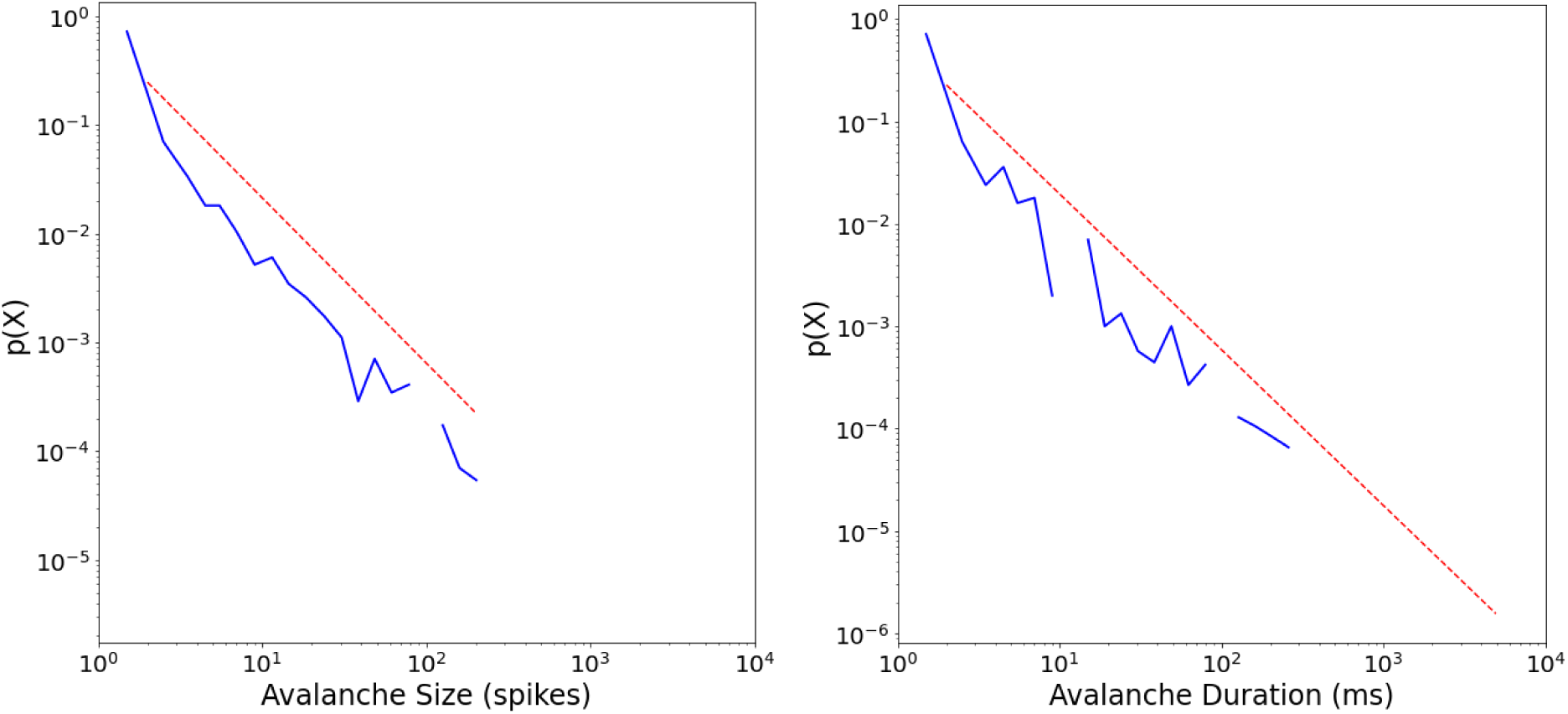
Power-law fits for avalanche size to all neurons in our 10 minute simulation of an M1 cortical column (sM1_11–22-2020_01). Left: Avalanche size probability density distribution with the number of avalanches normalized (y-axis) and the size of the avalanche (number of spikes; x-axis). Power-law fit equals −1.52 (red dashed line; sigma = 0.052, D = 0.023). Right: Avalanche duration probability density distribution showing the number of avalanches normalized (y-axis) and their durations (milliseconds; x-axis). Power-law fit equals −1.50 (red dashed line; sigma = 0.061, D = 0.038). Analyzed using the Python powerlaw package ^44^.

**Figure S8.**
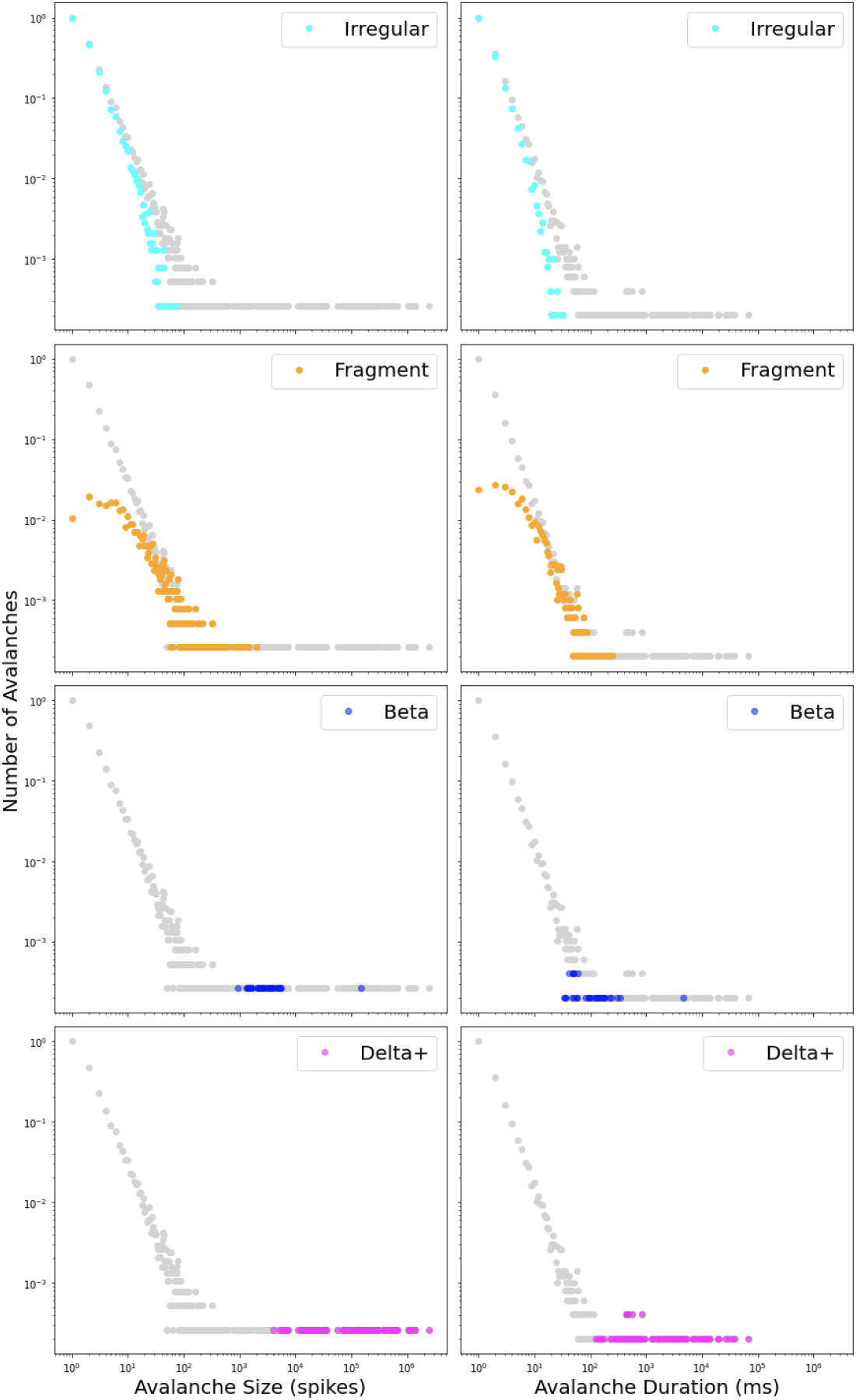
Log-log probability density distributions of avalanche size (first column) and duration (second column) for all avalanches (gray) and each of four avalanche types (sM1_11–22-2020_01; row 3; Fig S2).

**Figure S9.**
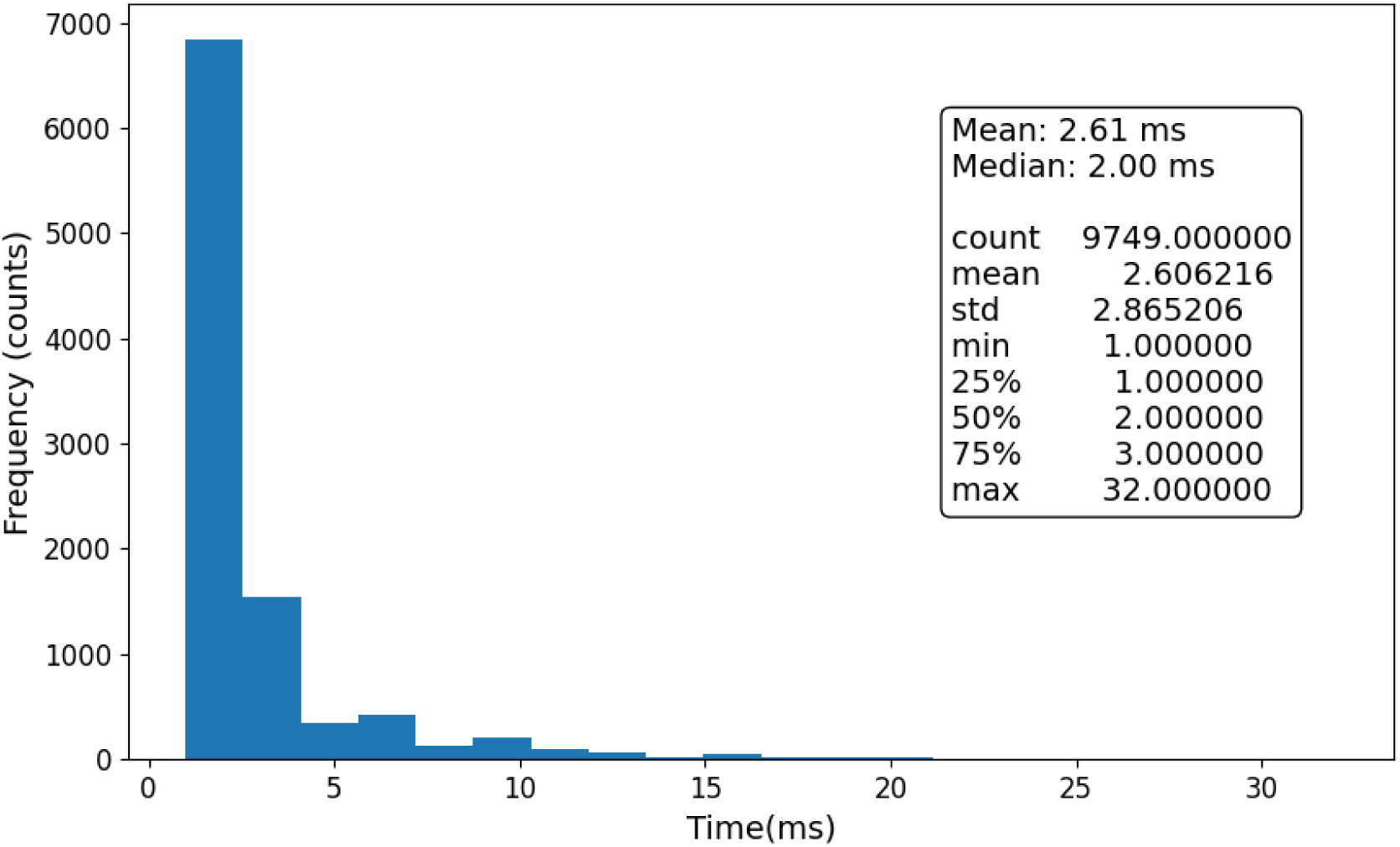
Inter-avalanche intervals in milliseconds (sM1_11–22-2020_01; row 3; Fig S2).

**Table S4.**
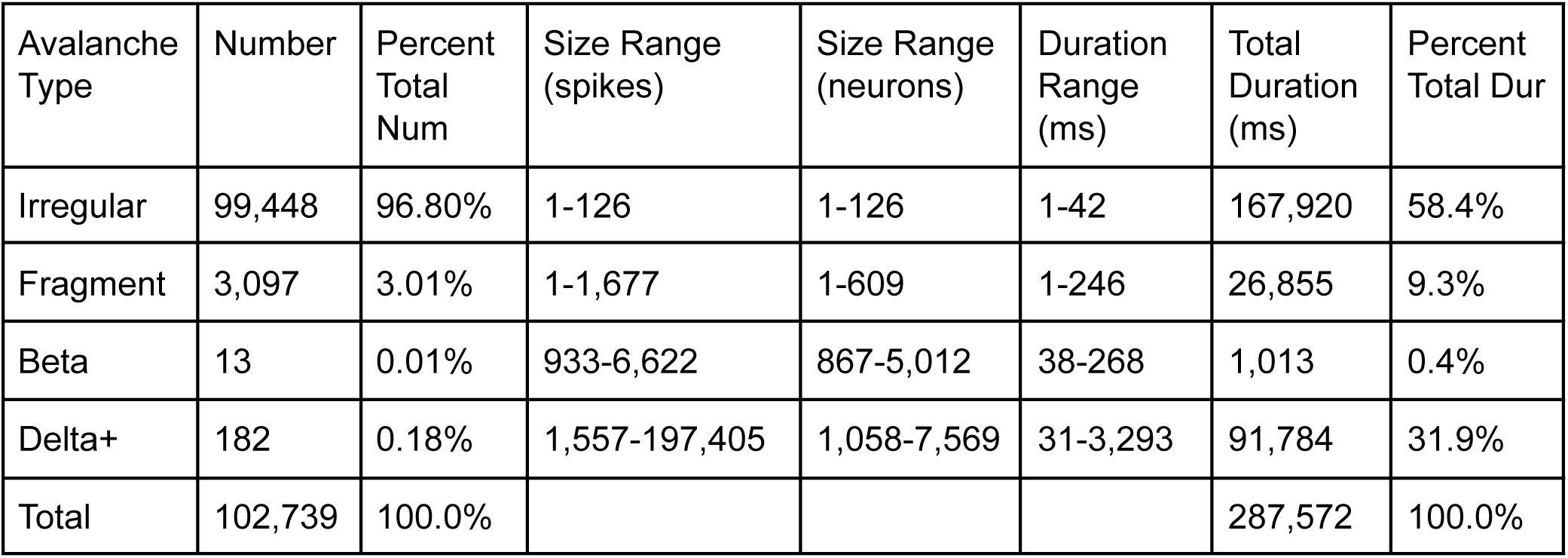
Data from a 10 minutes simulation of M1 (sM1_11-23-2020_05). Row 4 in Fig S2.

| Avalanche Type | Number | Percent Total Num | Size Range (spikes) | Size Range (neurons) | Duration Range (ms) | Total Duration (ms) | Percent Total Dur |
| --- | --- | --- | --- | --- | --- | --- | --- |
| Irregular | 99,448 | 96.80% | 1-126 | 1-126 | 1-42 | 167,920 | 58.4% |
| Fragment | 3,097 | 3.01% | 1-1,677 | 1-609 | 1-246 | 26,855 | 9.3% |
| Beta | 13 | 0.01% | 933-6,622 | 867-5,012 | 38-268 | 1,013 | 0.4% |
| Delta+ | 182 | 0.18% | 1,557-197,405 | 1,058-7,569 | 31-3,293 | 91,784 | 31.9% |
| Total | 102,739 | 100.0% |  |  |  | 287,572 | 100.0% |

**Figure S10.**
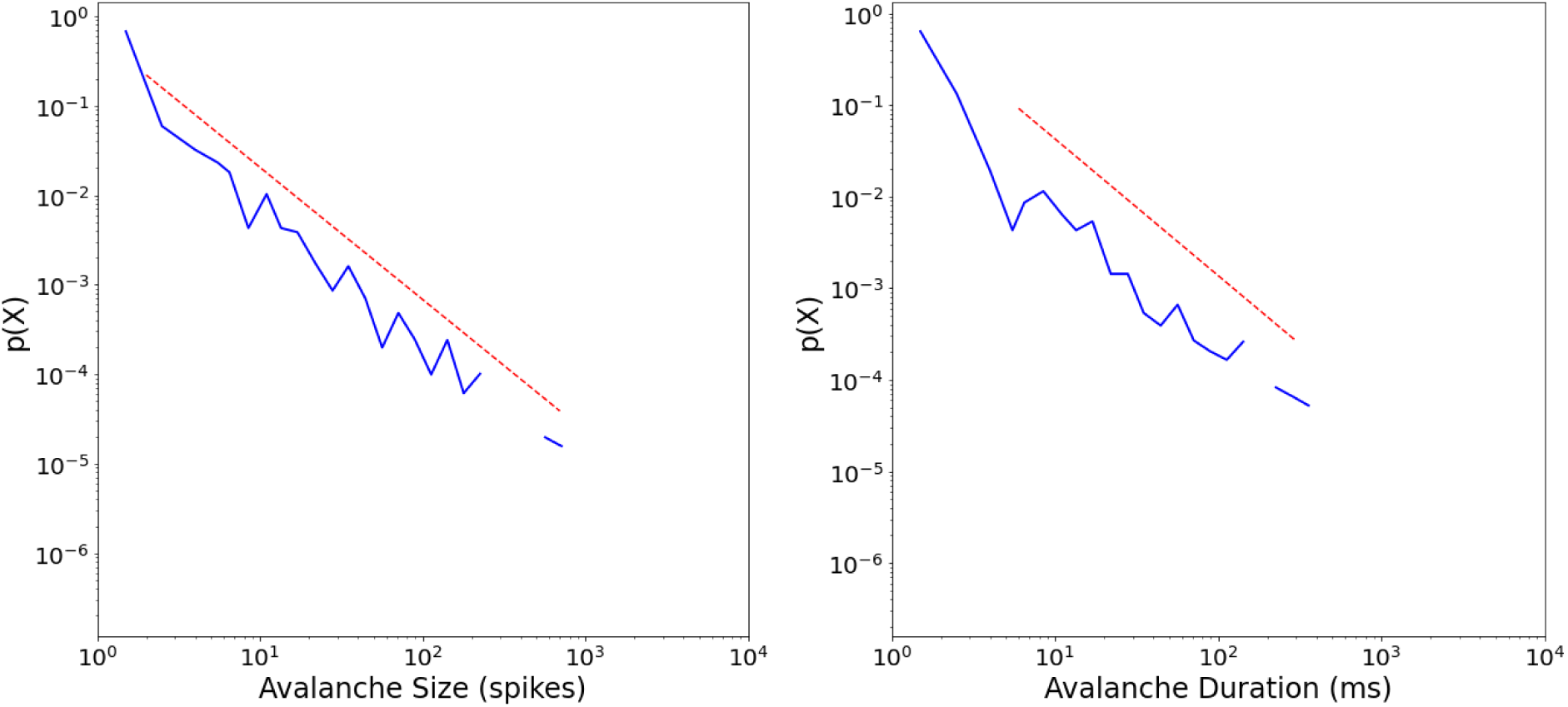
Power-law fits for avalanche size to all neurons in our 10 minute simulation of an M1 cortical column (sM1_11-23-2020_05). Left: Avalanche size probability density distribution with the number of avalanches normalized (y-axis) and the size of the avalanche (number of spikes; x-axis). Power-law fit equals −1.48 (red dashed line; sigma = 0.044, D = 0.028). Right: Avalanche duration probability density distribution showing the number of avalanches normalized (y-axis) and their durations (milliseconds; x-axis). Power-law fit equals −1.49 (red dashed line; sigma = 0.082, D = 0.057). Analyzed using the Python powerlaw package ^44^.

**Figure S11.**
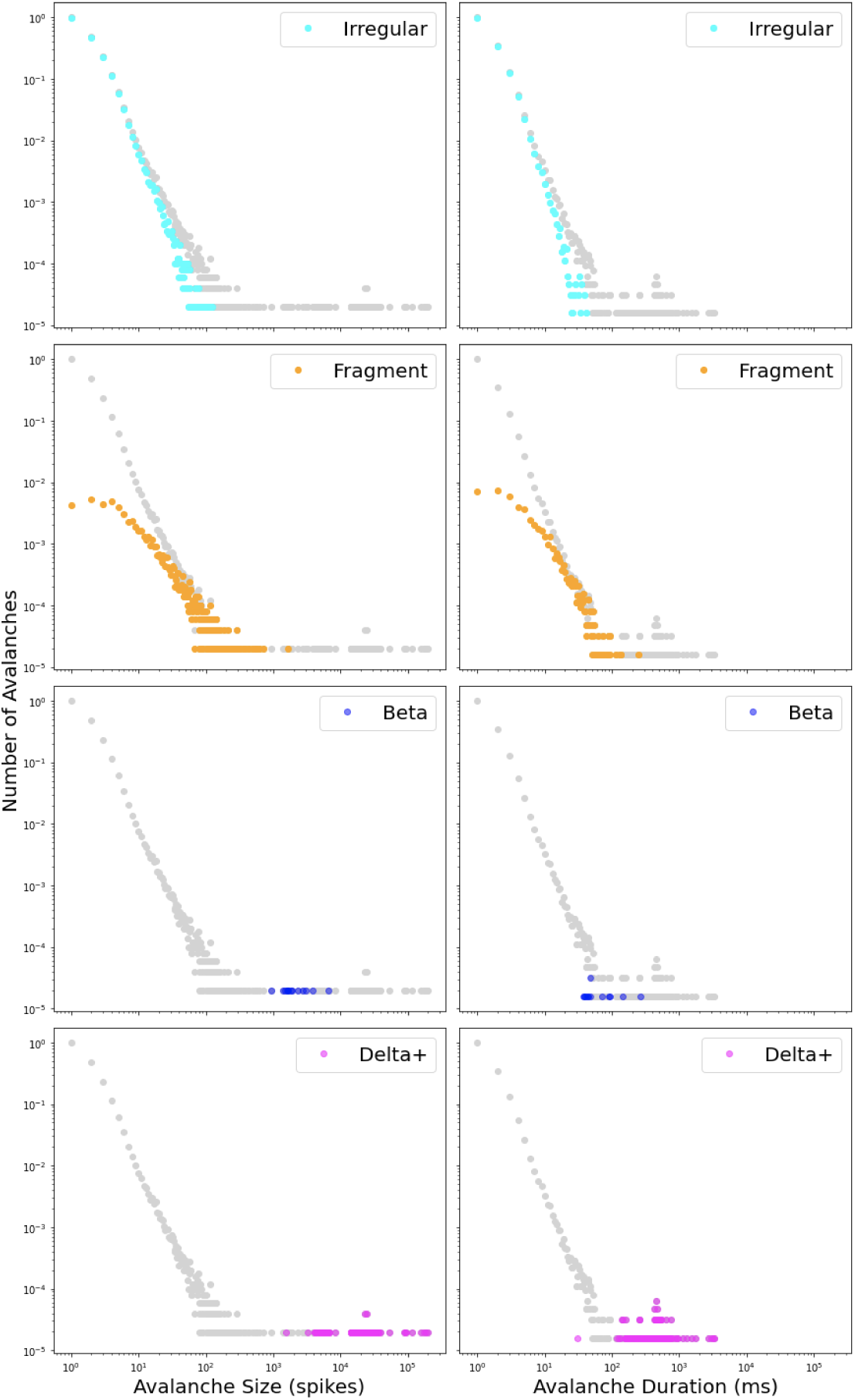
Log-log probability density distributions of avalanche size (first column) and duration (second column) for all avalanches (gray) and each of four avalanche types (sM1_11-23-2020_05; row 4; Fig S2).

**Figure S12.**
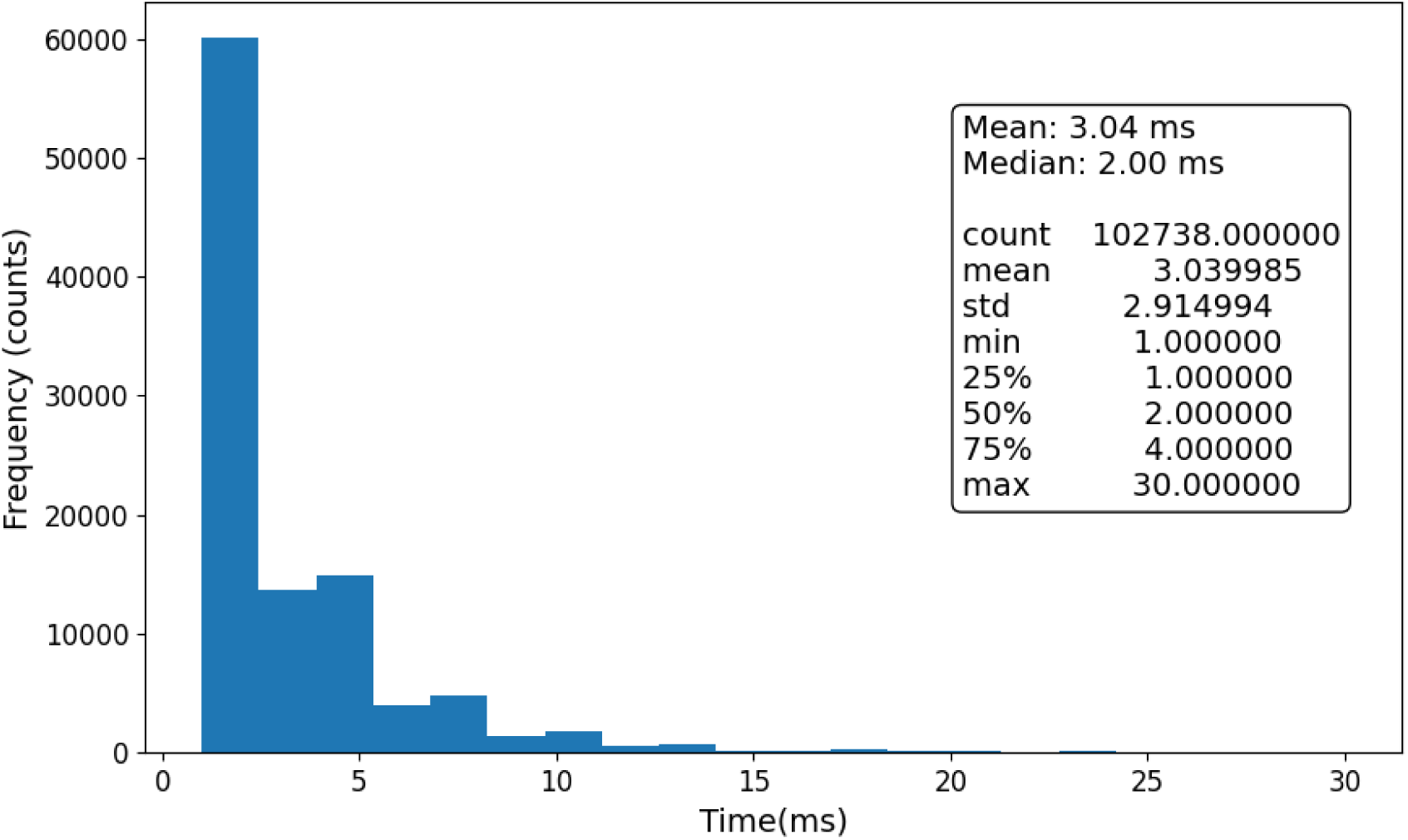
Inter-avalanche intervals in milliseconds (sM1_11-23-2020_05; row 4; Fig S2).

**Table S5.**
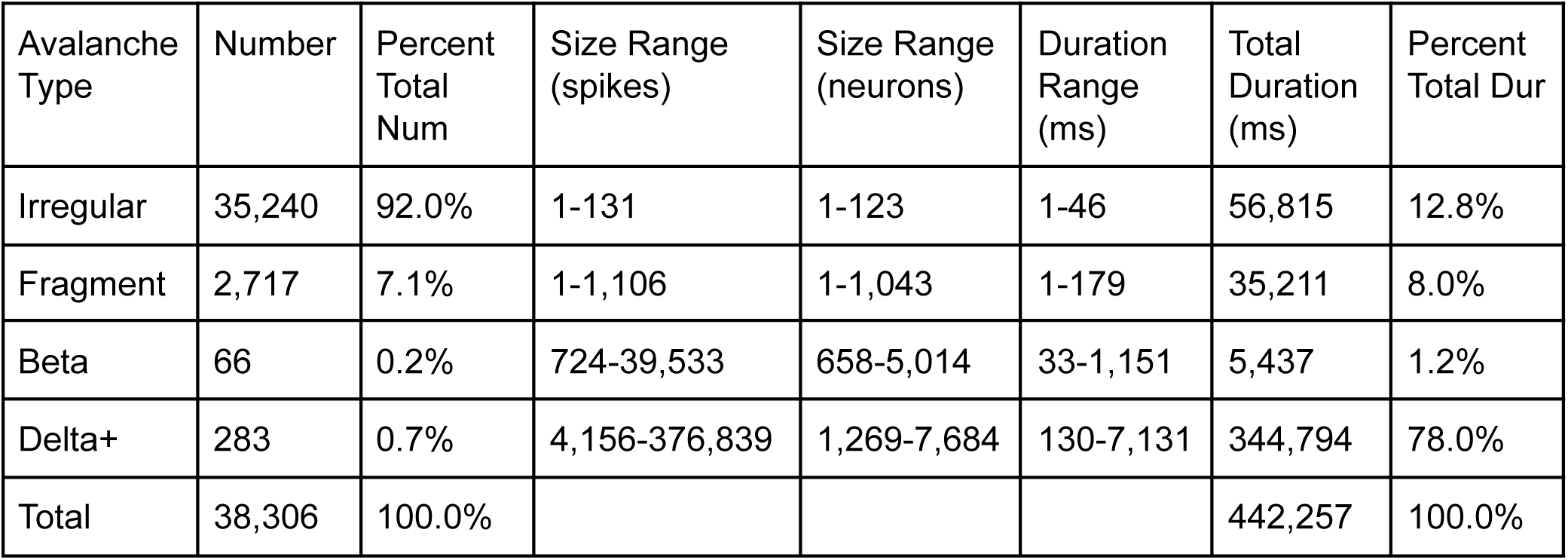
Data from a 10 minutes simulation of M1 (sM1_11–26-2020_01). Row 5 in Fig S2.

| Avalanche Type | Number | Percent Total Num | Size Range (spikes) | Size Range (neurons) | Duration Range (ms) | Total Duration (ms) | Percent Total Dur |
| --- | --- | --- | --- | --- | --- | --- | --- |
| Irregular | 35,240 | 92.0% | 1-131 | 1-123 | 1-46 | 56,815 | 12.8% |
| Fragment | 2,717 | 7.1% | 1-1,106 | 1-1,043 | 1-179 | 35,211 | 8.0% |
| Beta | 66 | 0.2% | 724-39,533 | 658-5,014 | 33-1,151 | 5,437 | 1.2% |
| Delta+ | 283 | 0.7% | 4,156-376,839 | 1,269-7,684 | 130-7,131 | 344,794 | 78.0% |
| Total | 38,306 | 100.0% |  |  |  | 442,257 | 100.0% |

**Figure S13.**
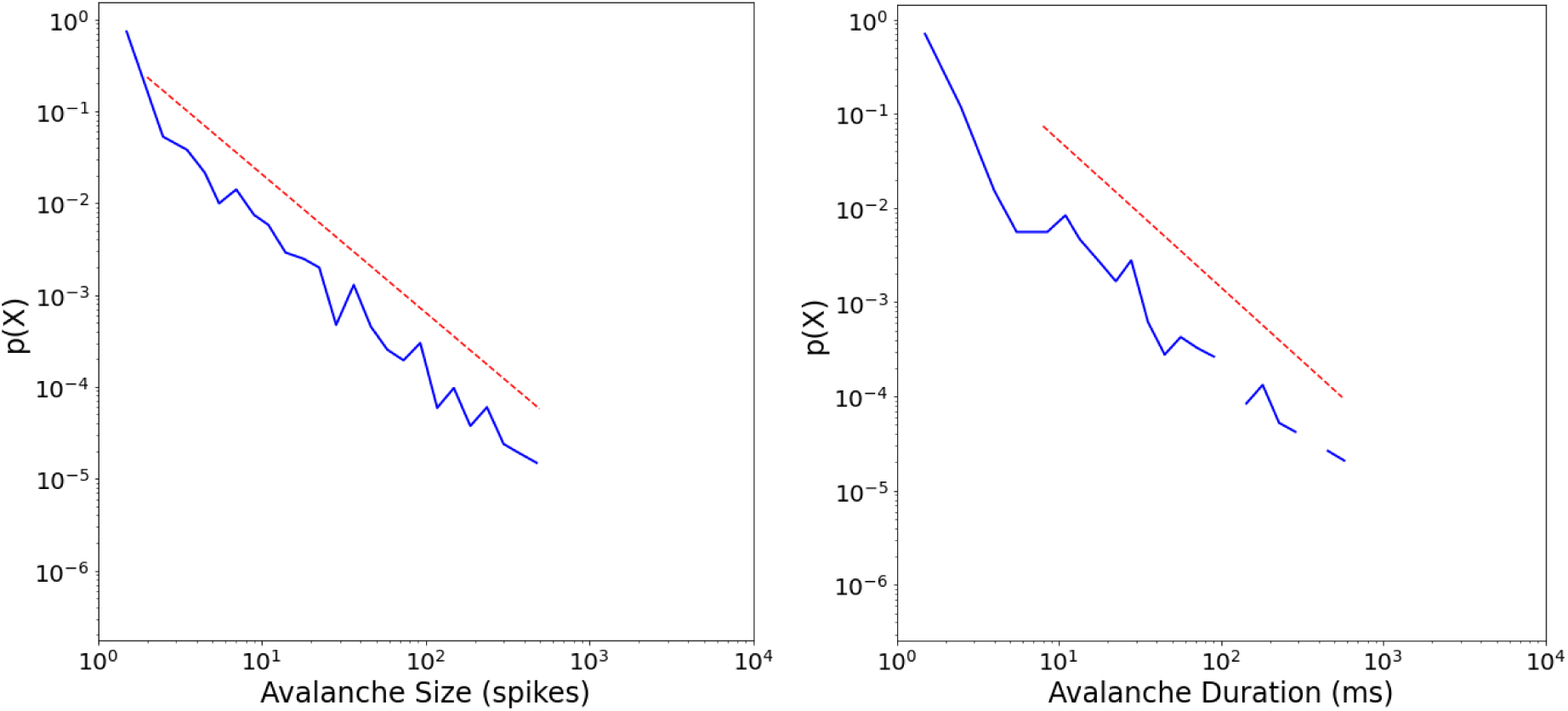
Power-law fits for avalanche size to all neurons in our 10 minute simulation of an M1 cortical column (sM1_11–26-2020_01). Left: Avalanche size probability density distribution with the number of avalanches normalized (y-axis) and the size of the avalanche (number of spikes; x-axis). Power-law fit equals −1.51 (red dashed line; sigma = 0.041, D = 0.045). Right: Avalanche duration probability density distribution showing the number of avalanches normalized (y-axis) and their durations (milliseconds; x-axis). Power-law fit equals −1.56 (red dashed line; sigma = 0.083, D = 0.057). Analyzed using the Python powerlaw package ^44^.

**Figure S14.**
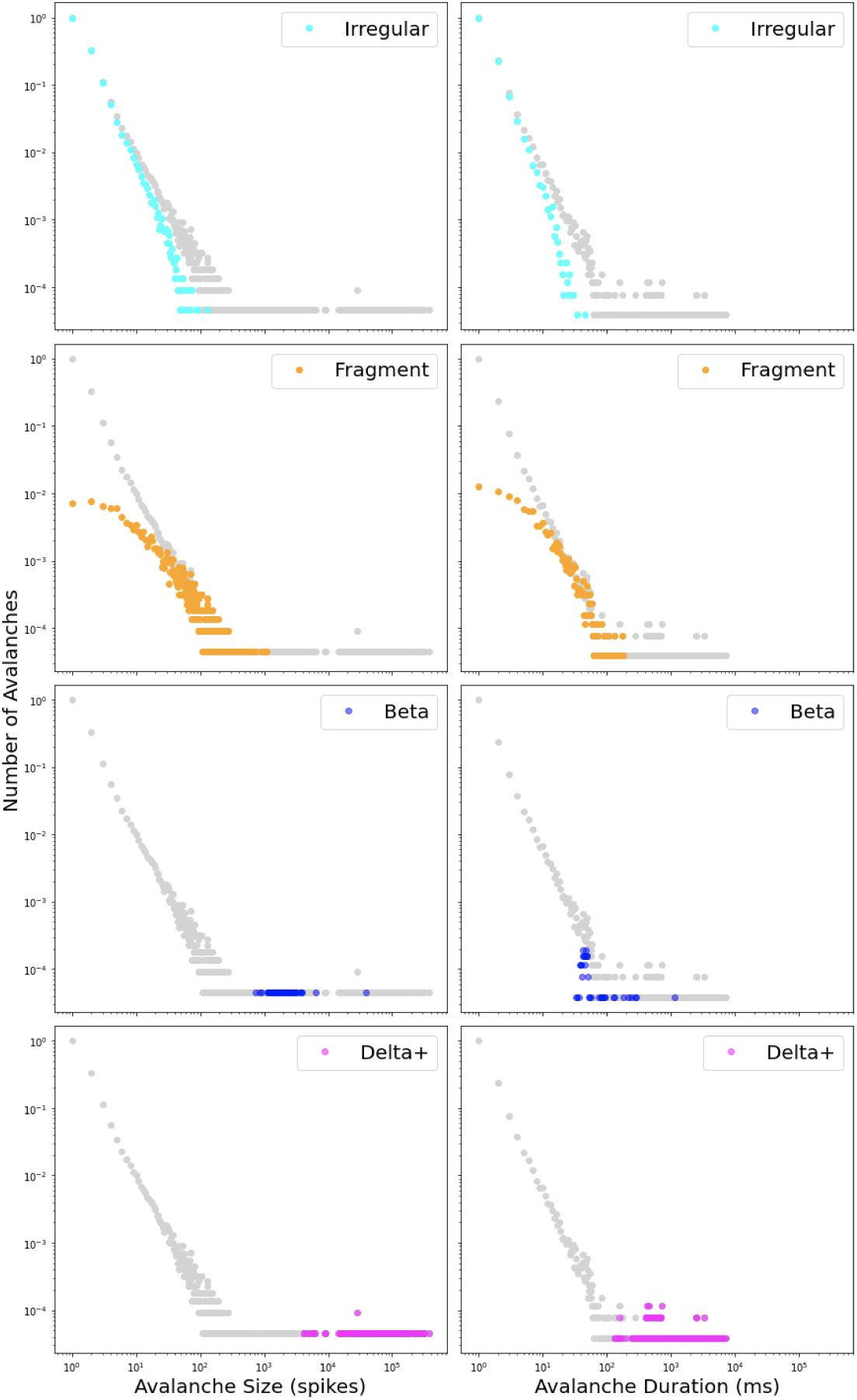
Log-log probability density distributions of avalanche size (first column) and duration (second column) for all avalanches (gray) and each of four avalanche types (sM1_11–26-2020_01; row 5; Fig S2).

**Figure S15.**
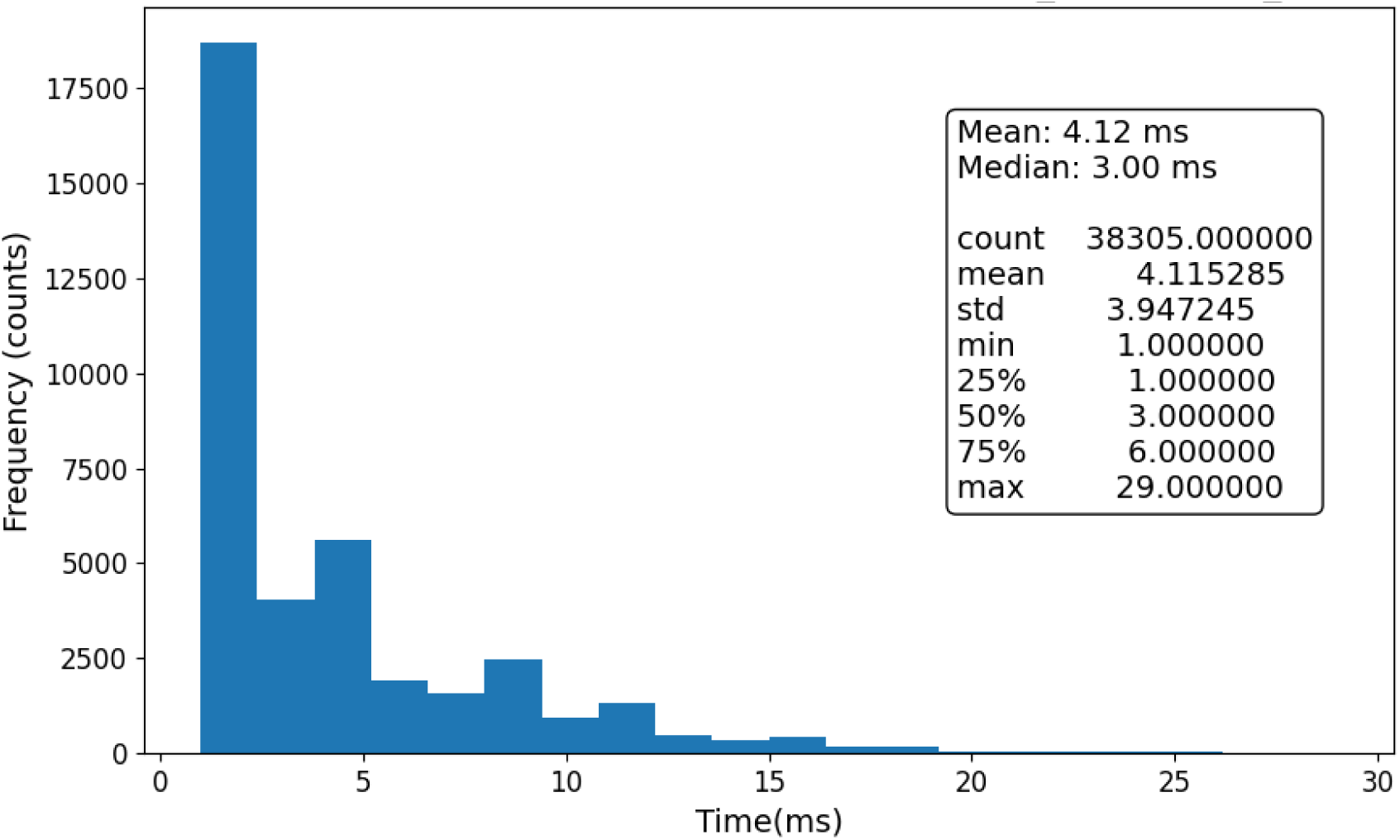
Inter-avalanche intervals in milliseconds (sM1_11–26-2020_01; row 5; Fig S2).

**Table S6.** Data from a 10 minutes simulation of M1 (sM1_11–28-2020_05). Row 6 in Fig S2.

| Avalanche Type | Number | Percent Total Num | Size Range (spikes) | Size Range (neurons) | Duration Range (ms) | Total Duration (ms) | Percent Total Dur |
| --- | --- | --- | --- | --- | --- | --- | --- |
| Irregular | 774 | 38.0% | 1-64 | 1-64 | 1-25 | 2599 | 0.4% |
| Fragment | 613 | 30.0% | 1-1,102 | 1-1,064 | 1-198 | 8615 | 1.4% |
| Beta | 259 | 12.7% | 554-5,692 | 492-4,338 | 31-310 | 15,218 | 2.6% |
| Delta+ | 393 | 19.3% | 4,335-352,729 | 1,307-7,719 | 101-8,681 | 570,323 | 95.6% |
| Total | 2,039 | 100.0% |  |  |  | 596,755 | 100.0% |

**Figure S16.**
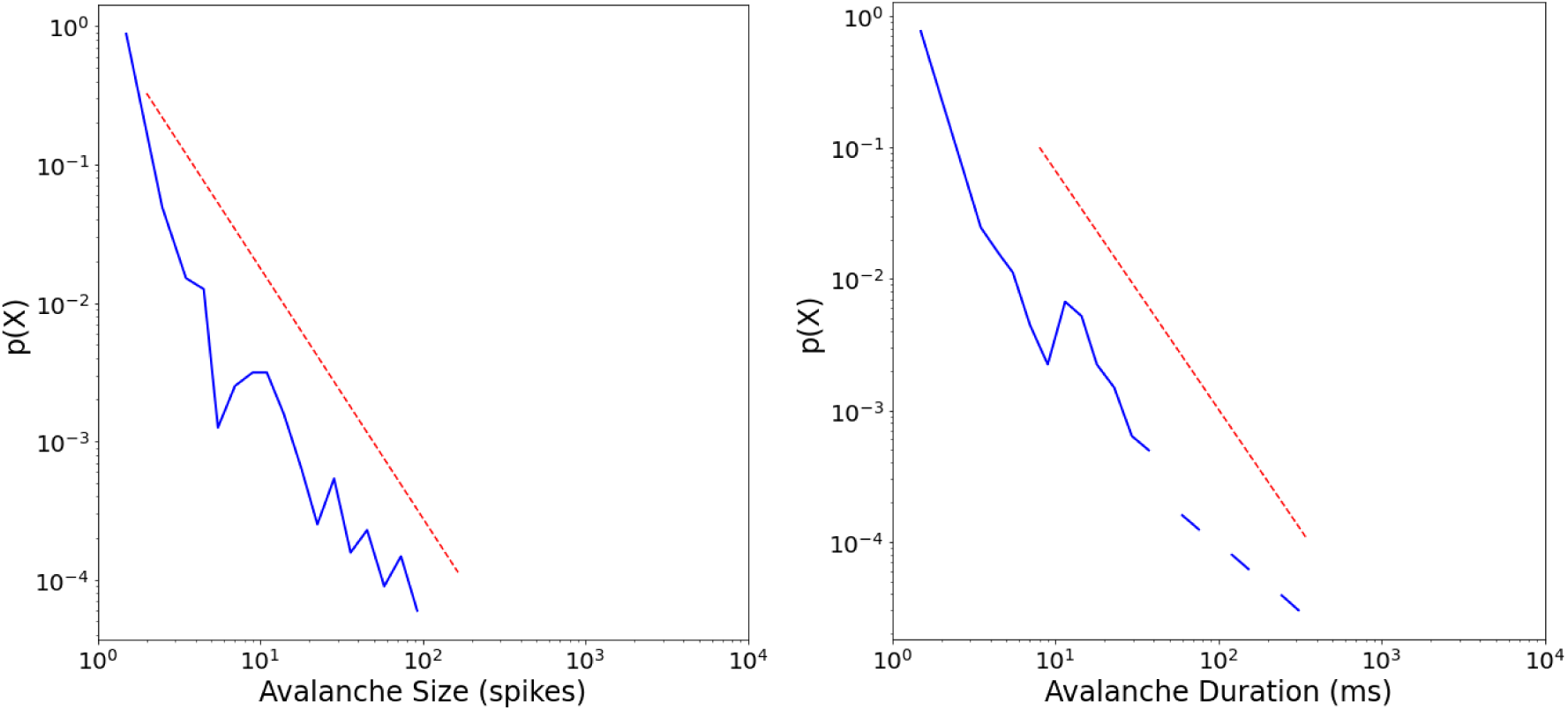
Power-law fits for avalanche size to all neurons in our 10 minute simulation of an M1 cortical column (sM1_11–28-2020_05). Left: Avalanche size probability density distribution with the number of avalanches normalized (y-axis) and the size of the avalanche (number of spikes; x-axis). Power-law fit equals −1.81 (red dashed line; sigma = 0.082, D = 0.075). Right: Avalanche duration probability density distribution showing the number of avalanches normalized (y-axis) and their durations (milliseconds; x-axis). Power-law fit equals −1.82 (red dashed line; sigma = 0.14, D = 0.13). Analyzed using the Python powerlaw package ^44^.

**Figure S17.**
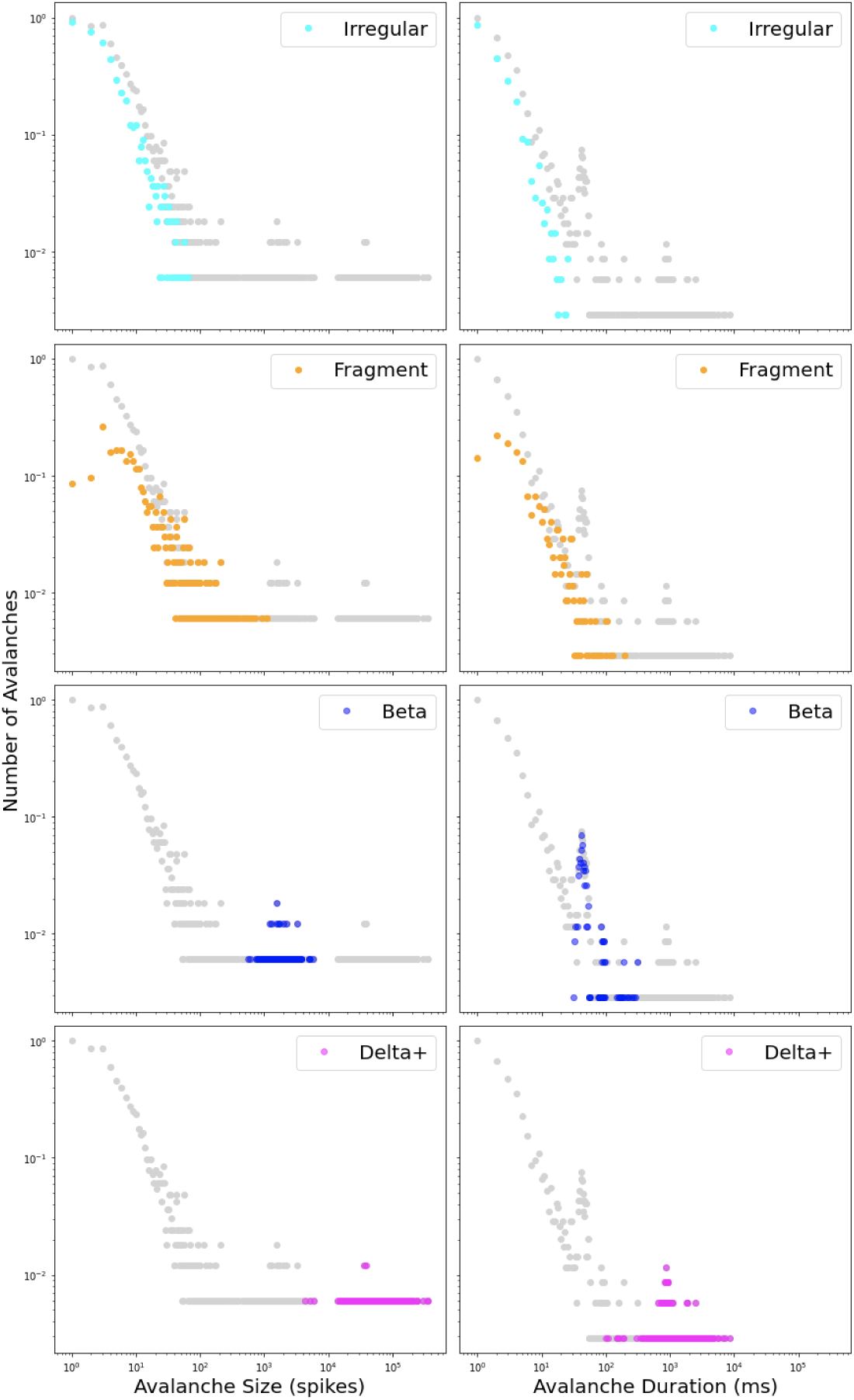
Log-log probability density distributions of avalanche size (first column) and duration (second column) for all avalanches (gray) and each of four avalanche types (sM1_11–28-2020_05; row 6; Fig S2).

**Figure S18.**
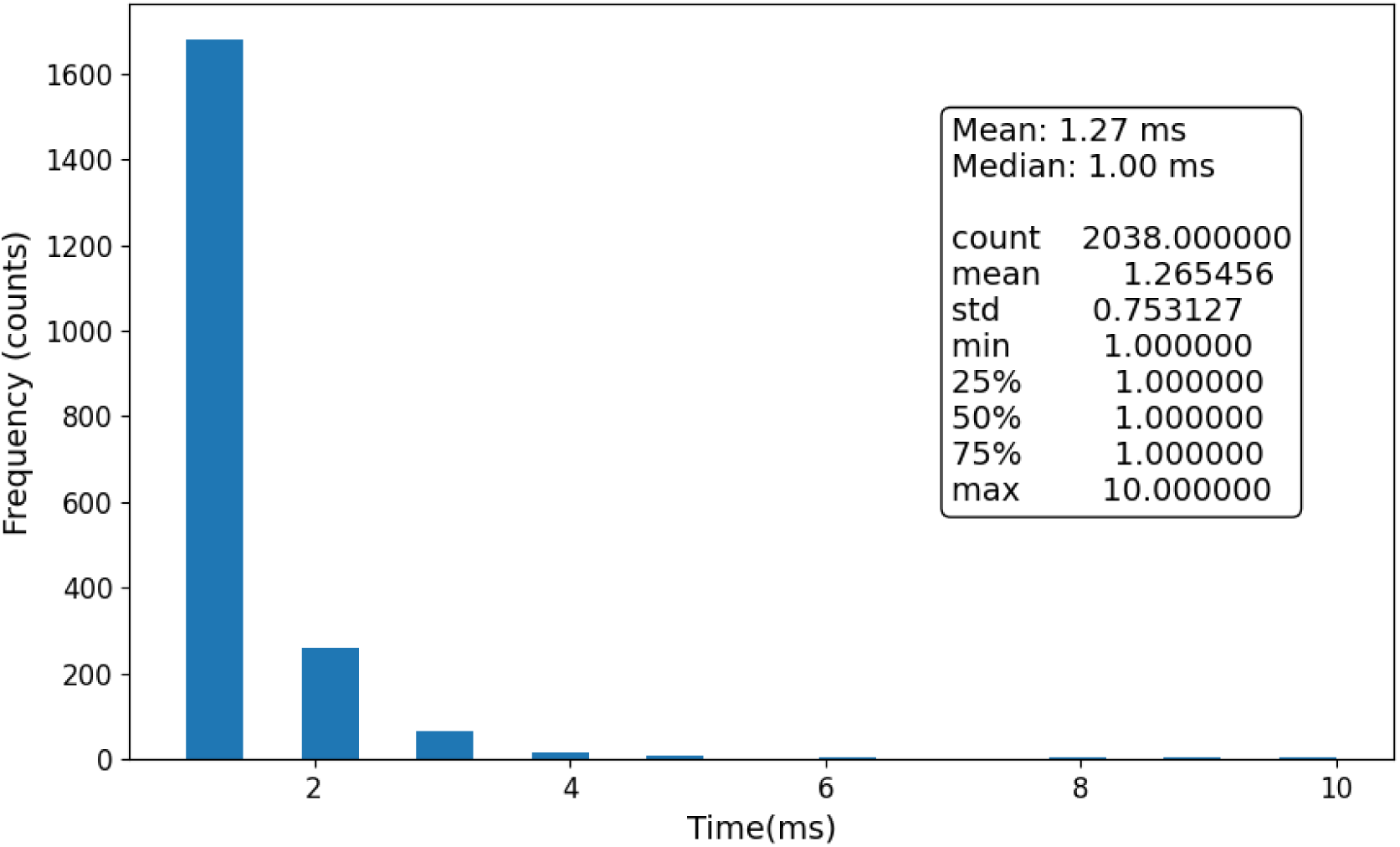
Inter-avalanche intervals in milliseconds (sM1_11–28-2020_05; row 6; Fig S2).

**Table S7.**
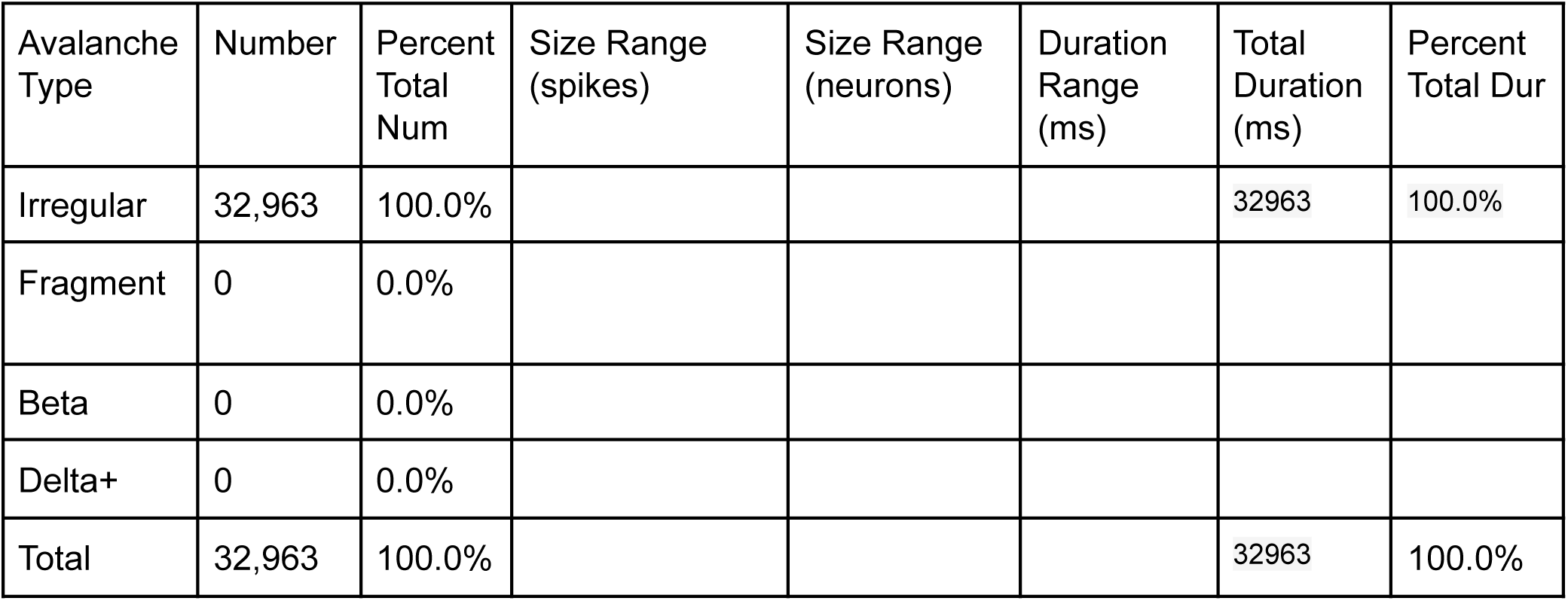
Data from a 10 minutes simulation of M1 (sM1_04–27-2021_17). Row 7 in Fig S2.

**Figure S19.**
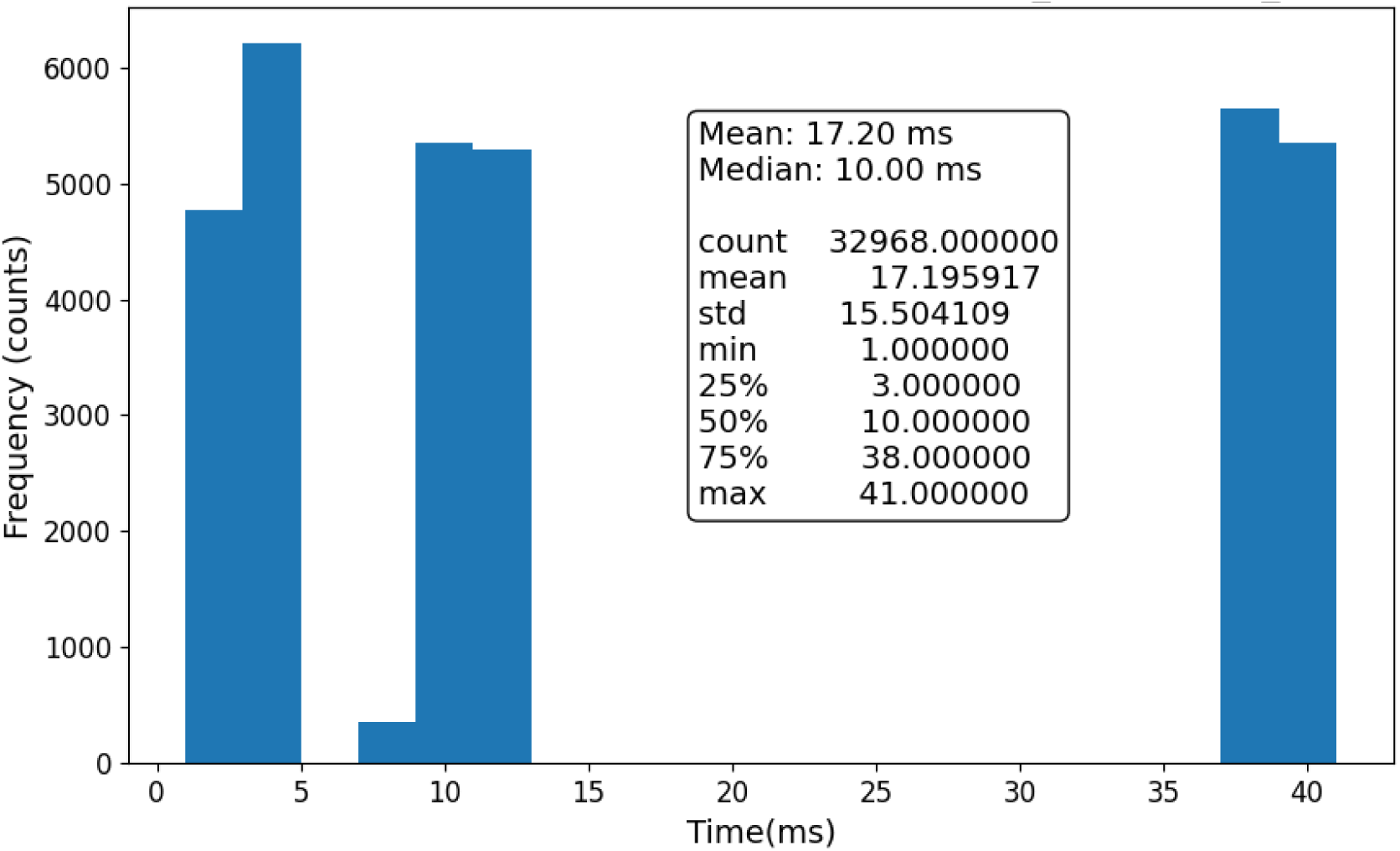
Inter-avalanche intervals in milliseconds (sM1_04–27-2021_17; row 7; Fig S2).

## Notes

### Competing Interest Statement

The authors have declared no competing interest.

